# Evaluating the Performance of the Astral Mass Analyzer for Quantitative Proteomics Using Data Independent Acquisition

**DOI:** 10.1101/2023.06.03.543570

**Authors:** Lilian R. Heil, Eugen Damoc, Tabiwang N. Arrey, Anna Pashkova, Eduard Denisov, Johannes Petzoldt, Amelia C. Peterson, Chris Hsu, Brian C. Searle, Nicholas Shulman, Michael Riffle, Brian Connolly, Brendan X. MacLean, Philip M. Remes, Michael W. Senko, Hamish I. Stewart, Christian Hock, Alexander A. Makarov, Daniel Hermanson, Vlad Zabrouskov, Christine C. Wu, Michael J. MacCoss

## Abstract

We evaluate the quantitative performance of the newly released Asymmetric Track Lossless (Astral) analyzer. Using data independent acquisition, the Thermo Scientific™ Orbitrap™ Astral™ mass spectrometer quantifies 5 times more peptides per unit time than state-of-the-art Thermo Scientific™ Orbitrap™ mass spectrometers, which have long been the gold standard for high resolution quantitative proteomics. Our results demonstrate that the Orbitrap Astral mass spectrometer can produce high quality quantitative measurements across a wide dynamic range. We also use a newly developed extra-cellular vesicle enrichment protocol to reach new depths of coverage in the plasma proteome, quantifying over 5,000 plasma proteins in a 60-minute gradient with the Orbitrap Astral mass spectrometer.

## Introduction

Quantitative proteomics has advanced significantly over the past decade, becoming a critical tool for biological and biomedical research. However, robust proteome-wide measurements remain challenging due to the complexity of the proteome. Notably, the dynamic range of the proteome is vast, with the cellular proteome spanning 7 orders of magnitude^1^ and some biofluids, such as plasma, spanning 10 orders of magnitude.^2^ Additionally, ultra-high sensitivity is more critical than ever as the field expands to include single cell measurements.^3–5^ Despite advances in technology, there is still room for improvement in quantitative sensitivity, accuracy, and precision across the complete dynamic range. While a large subset of well-characterized proteins are studied frequently, relatively little is known about many lower abundance proteins.^6^ Thus, mass spectrometry (MS) has emerged as the dominant method for quantitative proteome-wide measurements,^7^ however, further advancing the field will require continued improvement in hardware and data acquisition techniques.

Mass spectrometry offers the promise of comprehensive proteome-wide studies, but quantifying proteins accurately across the full dynamic range remains an unresolved challenge. Recent studies have reported the detection of large numbers of proteins: up to 7700 proteins in a 44-minute run^8^, 10,000 proteins in a single 120-minute run^9^, and over 12,000 proteins using biochemical fractionation.^10^ These results highlight the increasing difficulty in measuring low abundance proteins. Additionally, these studies each implement their own thresholds for false discovery rates and quantifiable peaks, making these results difficult to compare. Even a standard false discovery rate of 1% can have widely different meanings across different tools, instruments, and data sets. Measurements that do not produce any quantitative information inflate numbers while diluting statistical power. Yet, many proteomics studies take quantification for granted,^11^ with very few experiments to benchmark quantification over a significant dynamic range. Therefore, rigorous analytical evaluation of novel technologies, with a primary focus on their quantitative capabilities, is a critical part of the technology development cycle.

The Astral is a novel high resolution accurate mass (HRAM) analyzer that shares some of the operational principles with established mass analyzers, including the Orbitrap, ion trap and time-of-flight.^12^ A nearly lossless ion transfer is its most unique characteristic, derived from aligned ion injection and precise modulation of ion motion in three dimensions on a long asymmetric track and resulting in high sensitivity and consequently high analytical acquisition rate.^12^ Additionally, due to the configuration of the components (**Figure 1A**), automatic gain control (AGC)^13^ may be used to increase sensitivity and interspectrum dynamic range without the need for spectrum averaging as is the case with conventional orthogonal acceleration TOFs. The theoretical benefits of such an analyzer are evident, but the practical impacts still need to be assessed. To this end, we evaluated the quantitative performance of the Orbitrap Astral MS for data independent acquisition (DIA) and benchmark it relative to widely used Orbitrap based instruments.

**Figure 1.**
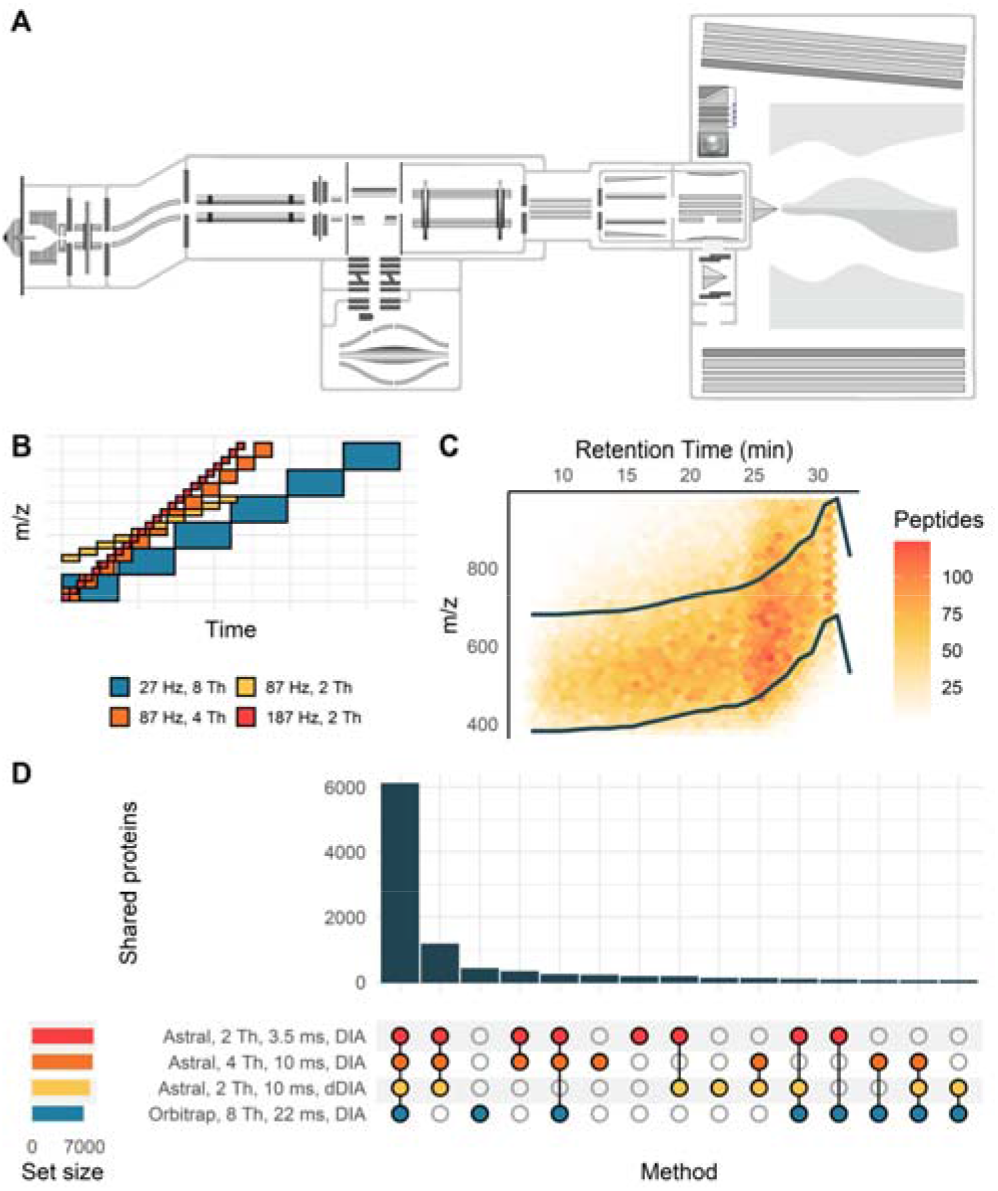
Quantitative comparison of the Orbitrap Astral MS and Orbitrap Fusion Lumos performance on a bulk cell digest. Schematic of the Orbitrap Astral mass spectrometer (A) and of isolation windows used for each method, with a limited mass range displayed (B). A dynamic DIA method was created based on peptide feature density with systematic isolation windows moving across the mass range between the two dark colored lines (C). The overlap of protein identifications between each method shows a high level of agreement between different Astral methods, with a subset of these proteins being detectable in the Orbitrap (D). Data were acquired on the Orbitrap Astral MS with a 24-minute gradient compared to a 90-minute gradient on the Orbitrap Fusion Lumos MS.

We use a matrix-matched calibration curve^11, 14^ to compare limits of quantification in both analyzers. This benchmark demonstrates the quantitative gains associated with the Orbitrap Astral mass spectrometer relative to existing state-of-the-art technology, with 22% more quantifiable peptides in a run that is about one-quarter of the length of that used in the Orbitrap. We further demonstrate quantitative gains associated with using dynamic DIA^15^ (dDIA) to maximize injection times across the mass range. To highlight the significance of this improved dynamic range, we analyze the plasma proteome. Using a membrane bound particle enrichment method,^16^ we quantified 5163 proteins in a single 70-minute LC-MS run with the Orbitrap Astral mass spectrometer. The benchmarking described here demonstrates the power of the Astral mass analyzer for robust quantitative data independent acquisition.

## Materials and Methods

### Cell Culture

HeLa S3 cells were cultured in DMEM/10% FBS, with stable isotope labeling performed according to manufacturer instructions using the Thermo Scientific™ SILAC™ Protein Quantitation Kit (Catalog A33972, Thermo Fisher Scientific). Briefly, cells were exchanged into DMEM (supplemented with Penicillin-Streptomycin and Thermo Scientific™ GlutaMAX ™ containing either unlabeled L-lysine and L-arginine, or the same supplemented DMEM containing 13C6 15N2 L-lysine and 13C6 15N4 L-arginine for SILAC labeling. Labeling was performed over 8 doublings and cells were then harvested at ∼85% confluence and pelleted by centrifugation.

### Matrix-matched calibration curve sample preparation

Both SILAC labeled and unlabeled HeLa cell lysates were prepared using protein aggregation capture.^17, 18^ Briefly, cell pellets were lysed with probe sonication in a lysis buffer containing 2% SDS. A Pierce BCA assay (Thermo Fisher Scientific) was used to estimate protein concentration and dilute lysates to a final concentration of 2 µg/µL in 1% SDS. Following reduction in 20 mM dithiothreitol and alkylation in 40 mM iodoacetamide, both samples were diluted to 70% acetonitrile and bound to MagReSyn Hydroxyl particles (ReSyn Biosciences) at a ratio of 100:1 (beads to protein). Subsequent washing was performed on a magnetic rack, with three washes in 95% acetonitrile followed by two washes in ethanol. Following the final wash, trypsin (in 50 mM ammonium bicarbonate) was added at an enzyme to protein ratio of 1:33. Samples were digested at 47° C for 4 hours, eluted from the beads and dried down via vacuum centrifugation. The digests were resuspended at a concentration of 200 ng/µL in 0.1% formic acid, Pierce peptide retention time calibration mix (PRTC) was spiked in to a final concentration of 5 pmol/µL, and digests were combined to form the following dilutions (X% unlabeled in (100-X)% heavy-labeled): 0, 0.5, 1, 5, 10, 25, 50, 70, 100%.

### Plasma preparation and membrane particle enrichment

An enriched membrane particle fraction was prepared from plasma using Mag-Net, a magnetic bead-based protocol developed by Wu et al.^16^ Plasma particle enrichment and protein aggregation capture^17, 18^ steps were performed on a Thermo Scientific™ KingFisher Flex.

Briefly, HALT cocktail (protease and phosphatase inhibitors, Thermo Fisher Scientific) was added to 100 µL plasma and then mixed 1:1 by volume with Binding Buffer (BB, 100 mM Bis-Tris Propane, pH 6.3, 150 mM NaCl). MagReSyn® strong anion exchange beads (ReSyn Biosciences) were first equilibrated 2 times in Equilibration/Wash Buffer (WB, 50 mM Bis Tris Propane, pH 6.5, 150 mM NaCl) with gentle agitation and then combined in a 1:4 ratio (volume beads:volume starting plasma) with the plasma:BB sample for 45 minutes at room temperature. The beads were washed with WB 3 times 5 minutes with gentle agitation. The enriched membrane particles on the beads were then solubilized and reduced in 50 mM Tris, pH 8.5/1% SDS/10 mM Tris (2-carboxyethyl) phosphine (TCEP) with 800 ng enolase standard added as a process control. Following reduction, the plate was removed from the Kingfisher Flex. Samples were alkylated with 15 mM iodoaceamide in the dark for 30 minutes and then quenched with 10 mM DTT for 15 minutes. Total unfractionated plasma (1 µL) was prepared in parallel (reduction, alkylation, and quenched) and added to the Kingfisher plate as a control for enrichment. The samples were processed using protein aggregation capture with minor modifications.^17, 18^ Briefly, the samples were adjusted to 70% acetonitrile, mixed, and then incubated for 10 minutes at room temperature to precipitate proteins onto the bead surface. The beads were washed 3 times in 95% acetonitrile and 2 times in 70% ethanol for 2.5 minutes each on magnet. Samples were digested for 1 hour at 47 °C in 100 mM ammonium bicarbonate with Thermo Scientific™ Pierce™ porcine trypsin at a ratio of 20:1 trypsin to protein. The digestion was quenched in 0.5% formic acid and spiked with Pierce Retention Time Calibrant peptide cocktail (Thermo Fisher Scientific) to a final concentration of 50 fmol/µL. Peptide digests were lyophilized and stored at -80°C.

### Liquid Chromatography-Mass spectrometry

All data were acquired using a Thermo Scientific™ Vanquish™ Neo UHPLC system coupled to an Orbitrap Astral mass spectrometer with a Thermo Scientific™ Easy-spray™ source. Data were acquired in data independent acquisition mode with a normalized collision energy of 25% and a default charge state of 2. A 110 cm Thermo Scientific™ µPac™ Neo HPLC column was used for the quantitative experiments with a 24-minute gradient of from 4 to 45% B and a flow rate of 750 nL/min. For plasma experiments, a two step 30 or 60 minute gradient at 750 nL/min from 4 to 45% B was used on a 110 cm µPac Neo HPLC column. Mobile phase A was 0.1% formic acid in water, and mobile phase B was 80% acetonitrile and 0.1% formic acid in water. For Orbitrap experiments, MS1 spectra were acquired in the Orbitrap every 0.6 s at a resolution of 30,000, and MS/MS spectra were acquired in the Orbitrap at 15,000 resolving power with a maximum injection time of 23 ms. For the Orbitrap Astral MS experiments, MS1 spectra were acquired in the Orbitrap at a resolving power of 240,000 every 0.6 s, and MS/MS spectra were acquired in the Astral analyzer with varying injection times noted for each experiment (**Supplemental Table 1)**. In all MS/MS experiments, DIA precursor isolation windows spanned 380-980 Th unless noted otherwise. The MS1 mass range was the same as the MS/MS precursor range. All experiments besides chromatogram libraries were acquired in triplicate.

A number of MS/MS parameters were used and are summarized in **Supplemental Table 1.** For the chromatogram library runs, the same parameters were used in both the Orbitrap and the Astral, with 6 gas phase fractions each spanning a 100-Da range with 4 Da isolation windows and a 23 ms maximum injection time. For the dilution curve, data were acquired from low concentration to high concentration, completing one replicate from each method before running a blank and starting the second replicate at the lowest concentration. Raw data files, EncyclopeDIA search results, and all Skyline documents have been deposited in the ProteomeXChange Consortium (identifier PXD042704) via Panorama Public (https://panoramaweb.org/AstralBenchmarking.url).

A reference dataset from Heil et al, 2023^15^ acquired with the Orbitrap was used. Briefly, the dataset was acquired using the Orbitrap mass analyzer on a Thermo Scientific™ Orbitrap Fusion Lumos™. Separation was performed with a 30 cm packed column attached to a Thermo Scientific™ EASY-nano™ LC across a two-step 90-minute gradient from 0 to 75% B at a flow rate of 300 nL/min. Mobile phase A was 0.1% formic acid in water, and mobile phase B was 80% acetonitrile and 0.1% formic acid in water. MS/MS spectra were acquired in the Orbitrap at 15k resolving power and a 22 ms maximum injection time using 8 Th isolation windows spanning a mass range from 400-1000 Th.

### Dynamic data independent acquisition method development

A single injection on Astral was analyzed with EncyclopeDIA using the Pan-Human library^19^ to create a preliminary list of peptides in the sample. Then, dynamic DIA boundaries were selected as described by Heil et al. (2023)^15^ to maximize the number of peptides covered in a 300 Th mass range with either 2 or 4 Th isolation windows. In dynamic DIA, the 300 Th mass range that contains the highest peptide feature density is divided into equal isolation windows. This mass range is adjusted across time while the number and size of isolation windows remains the same (**Figure 1C).**

### Data analysis

Prior to searching, raw files were converted to mzMLs using Proteowizard MSConvert (from ProteoWizard release 3.0.22335).^20^ For each experiment, a chromatogram library was generated by searching the 6-fraction chromatogram library runs against a Prosit^21^ predicted spectral library in EncyclopeDIA (v 3.0.0).^22^ Then, the quantitative DIA runs were searched against the appropriate chromatogram library in EncyclopeDIA with V2 scoring enabled and adjust inferred retention time boundaries set to true, and the quantitative results were loaded into Skyline (version 22.2.1.335)^23^ for protein grouping and subsequent analysis. Orbitrap data were searched and integrated with a 10 ppm mass tolerance, Astral data were searched with 10 ppm mass tolerance and integrated with 25 ppm mass tolerance to account for space charging at the apex of the peak. Conversion to mzML and importing into Skyline were done using the ProteoWizard Docker image available on Docker Hub at “ proteowizard/pwiz-skyline-i-agree-to-the-vendor-licenses:3.0.22335-b595b19”. Encyclopedia was run in a Docker image available Docker Hub at “mriffle/encyclopedia:3.0.0-rc.3”.

For the HeLa dilution curve, all 27 quantitative runs for a given method (3x 9 dilution points) were searched together in EncyclopeDIA and integrated in Skyline without any normalization. A version of Skyline that can refine transitions to give the best lower limit of quantification. This feature will be available in future Skyline releases. Peak areas and lower limits of quantification were exported from Skyline for further analysis.

For plasma data, all six (3x total plasma and 3x enriched plasma) were searched together in EncyclopeDIA and imported into Skyline for quantification. TIC normalized peak areas were exported from Skyline and used for subsequent analysis.

The downstream analysis code used to generate figures can be found on GitHub at: https://github.com/uw-maccosslab/AstralBenchmarking.

## Results and Discussion

### Peptide detections in a whole cell lysate

To evaluate the performance of the Astral analyzer, we performed a number of comparisons with the Orbitrap. First, we used gas phase fractionation^22, 24, 25^ to generate in-depth chromatogram libraries to compare the total peptide and protein level coverage with each analyzer in the same 24-minute gradient. We use these libraries for data analysis but they are also a way to compare deep proteome coverage when acquisition time is not a limiting factor. The philosophy of our workflow is that the chromatogram library should detect a superset of all observable peptides, while subsequent acquisitions may collect lower quality data, albeit at a faster rate. Here, we created a library from six different injections of the same HeLa digest with each analyzer, each covering a 100 Th mass range with 4 Th isolation windows and a 23 ms maximum injection time. Although the Astral has the ability to acquire spectra at rates up to 200 Hz, for this comparison we reduced the rate significantly and allowed a relatively long maximum injection time so that we could directly compare coverage with the Orbitrap. Even controlling for injection times between analyzers, we can expect to see differences because the Orbitrap fills with significantly more charges than the Astral (**Figure S1A**). While both mass analyzers tended to reach their maximum injection time in every spectrum (**Figure S1B**), the high sensitivity of the Astral means it has more data points per spectrum (**Figure S1C**) and decreased overhead enabled by additional parallelization (i.e. processing 5 distinct ion packets in parallel) allows the Astral to acquire spectra at a faster rate even at the same injection time (**Figure S1D**). As a result of these differences, there are 70% more detectable peptides in the Astral chromatogram library than the Orbitrap-based chromatogram library (87,257 and 51,603 respectively). Although it should be noted that these Orbitrap parameters may have been suboptimal and longer injection times could have resulted in deeper coverage.

In contrast to gas-phase fractionation approaches to build a library, the majority of DIA experiments performed today are single runs for quantitative analysis. To evaluate the ability of the Astral to detect peptides across a wide dynamic range, we acquired DIA data for a whole cell HeLa lysate in the Astral analyzer and compared the results to an existing dataset acquired by Heil et al. (2023) on the Orbitrap analyzer in an Orbitrap Fusion Lumos mass spectrometer.^15^ There are several differences between these datasets, most notably a 24-minute gradient at 750 nL/minute was used for the Astral data while the Orbitrap data was collected with a 90-minute gradient at 300 nL/minute. The longer gradient better accommodates the Orbitrap acquisition rate and should give it a significant advantage in this head-to-head comparison. Narrow window chromatogram libraries^22^ were generated for each method and used for analysis of the quantitative data, using various acquisition schemes in the Astral to compare with the Orbitrap (**Figure 1B, Supplemental Table 1**). Although sample specific libraries may not be necessary for the Astral when narrow (2-4 Th) isolation windows can be used across the full mass range, we use the narrow window library here for consistency and to increase search speed.

When creating a quantitative DIA method it is important to strike a balance between cycle time, spectrum acquisition rate, and isolation window width that fits the specific sample type, experimental goals, and LC parameters. Therefore, we evaluated three different Astral methods against a standard Orbitrap method. The first Astral method was acquired at 187 Hz using 3.5 ms maximum injection times and 2 Th isolation windows. The second Astral method used longer injection times (10 ms) and wider isolation windows (4 Th) in an effort to increase sensitivity and dynamic range. The third method used a dynamic DIA scheme^15^ which adjusts the isolation window range across time (**Figure 1C),** allowing for longer injection times (10 ms) with narrow isolation windows (2 Th). Generally, longer injection times allowed the Astral to accumulate more ions (**Figure S2A**) while slowing the overall acquisition rate significantly (**Figure S2B&D**). Additionally, narrower isolation windows reduced the spectral complexity, although spectra generated by the Astral are significantly more complex than those of the Orbitrap (**Figure S2C**). In this 24-minute gradient, all three Astral methods identified 7700-8200 proteins (**Figure 1D**) and 58,000 - 75,000 peptides (**Figure S3, Supplemental Table 2**). The Orbitrap detected 6885 proteins and 55,981 peptides in a 90-minute gradient, meaning that the Astral analyzer is able to detect more peptides and proteins in a 3.5x - shorter time than the Orbitrap analyzer.

To test performance of the Orbitrap while controlling the isolation window and using the same LC setup, we acquired a set of runs in the Orbitrap covering a 75 Th mass range with 2 Thisolation windows. As we observed with the 90-minute gradient, spectra generated in the Orbitrap typically consist of more ion signal distributed across fewer fragment ion peaks, pointing to the increased sensitivity of the Astral analyzer (**Figure S2**). In the same mass range, which was a subset of the total mass range covered, the Astral analyzer detected 43% more peptides (**Figure S4**). This observation suggests that part, but not all, of the increase in peptide detections seen in the Astral analyzer can be attributed to its high speed and sensitivity allowing it to cover a wider mass range with relatively narrow isolation windows.

### Quantitative benchmarking of DIA performance

While peptide and protein level identifications can be useful metrics to evaluate instrument performance, these numbers may not fully reflect the quality of the data. Therefore, we evaluated the quantitative precision and accuracy of the Orbitrap Astral MS using a 24-minute gradient relative to the Orbitrap Fusion Lumos using a 90-minute gradient. First, we looked at the technical precision across injection replicates with different inputs (**Figure 2**). We found that the coefficient of variation for 1 µg, 500 ng, and 100 ng of HeLa digest tends to be similar if not slightly better for the Astral than the Orbitrap. When compared with the same gradient and reduced mass range in the Orbitrap, a similar trend can be observed, suggesting that these results were not just due to differences in the chromatography or experimental setup (**Figure S5**). Additionally, we observed that using longer maximum injection times in the Astral analyzer improved the technical precision, with the narrow windows and longer injection times afforded by dynamic DIA providing an improvement in technical precision at all input levels.

**Figure 2.**
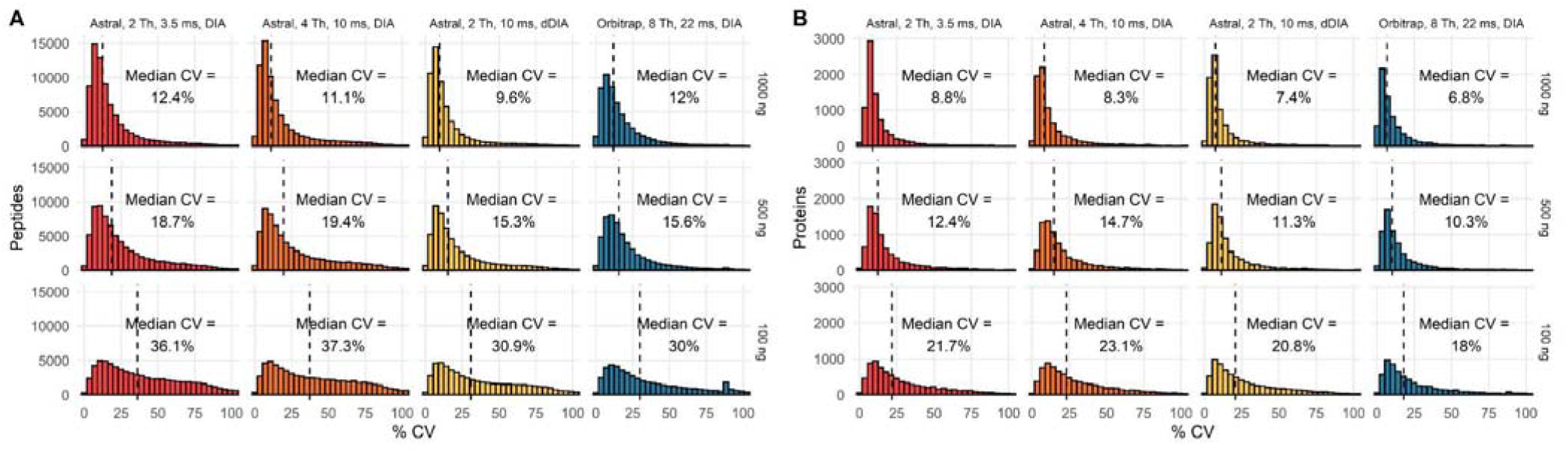
Evaluation of the technical precision of the data obtained with the Astral analyzer. Peptide (A) and protein (B) level coefficients of variation across 4 different acquisition methods. CVs are plotted for the same set of peptides with different amounts of HeLa injected with SILAC HeLa added as a background to keep total protein loading level constant (1000 ng). Dashed horizontal line represents the median CV, which is also indicated on the plot.

It is notable that the Astral analyzer is able to measure more peptides than the Orbitrap analyzer and still have similar if not better median CVs. Generally, the variance will increase as analyte abundance decreases. However, these results suggest that the Astral is able to generate very high-quality measurements for a large number of peptides in a short amount of time.

Technical precision is an important metric for evaluating performance, but quantitative accuracy is the goal of most proteomics experiments and needs to be evaluated independently. Therefore, we use matrix-matched calibration curves to evaluate proteome-wide quantitative performance.^11, 14^ By diluting our analyte (HeLa cell digest) in a similar matrix (SILAC labeled HeLa cell digest), we calculate the lower limits of quantification for thousands of peptides at a time and assign a robust analytical figure of merit to sensitivity. We can categorize peptides as quantitative if they can be assigned a lower limit of quantification (LLOQ), meaning that their signal has a linear response to changes in input when the total protein input is less than 1 µg. This measure provides more information than variance-based measurements, which can be misleading, as the variance of background signals can be quite low. This comparison is especially important here, where the mass analyzers have drastically different levels of background signal.

Using a 9-point matrix-matched calibration curve, we quantified at most 46,038 peptides in the Astral in a 24-minute gradient compared to 40,098 peptides in the Orbitrap in a 90-minute gradient (**Figure 3A, Supplemental Table 2**). The 9-point curve is an abbreviated version of the 13-point curves reported by Pino et al.^14^ to allow for the testing of more methods with limited instrument time. In addition to quantifying more peptides than the Orbitrap, the Astral tends to produce slightly better LLOQs than the Orbitrap for the peptides that both analyzers can quantify (**Figure 3B**). We can further improve quantification by integrating only the set of fragment ions that yield the best lower limit of quantification per peptide.^26^ This procedure improves the lower limit of quantification in all cases but is particularly beneficial for the Astral-based data, likely due to its higher spectral complexity (**Figure 3A&C**). With the refined transition set, the Astral quantifies at most 58,330 peptides (17,454 across at least a 10x linear dynamic range) while the Orbitrap quantifies 47,671 peptides (10,686 across at least a 10x linear range). In all cases, the Astral quantifies more peptides and tends to have a better LLOQ (**Figure 3C**). This quantification is achieved in significantly less time in the Astral than the Orbitrap. In fact, per unit time, the Astral quantifies 5 times as many peptides as the Orbitrap (**Figure S6**). Another way to look at the quantitative performance is to observe the ratio of signals between two dilution points. There, the sensitivity of the Astral is evident over a 10x dilution, where accurate quantification is observed for most peptides and proteins (**Figure 3D, Figure S7**). To eliminate the possibility that experimental differences were responsible for the increase in sensitivity, we also confirmed that the Orbitrap underperformed relative to the Astral in the reduced mass range 24-minute gradients, with considerably less sensitivity by all metrics (**Figure S8**).

**Figure 3.**
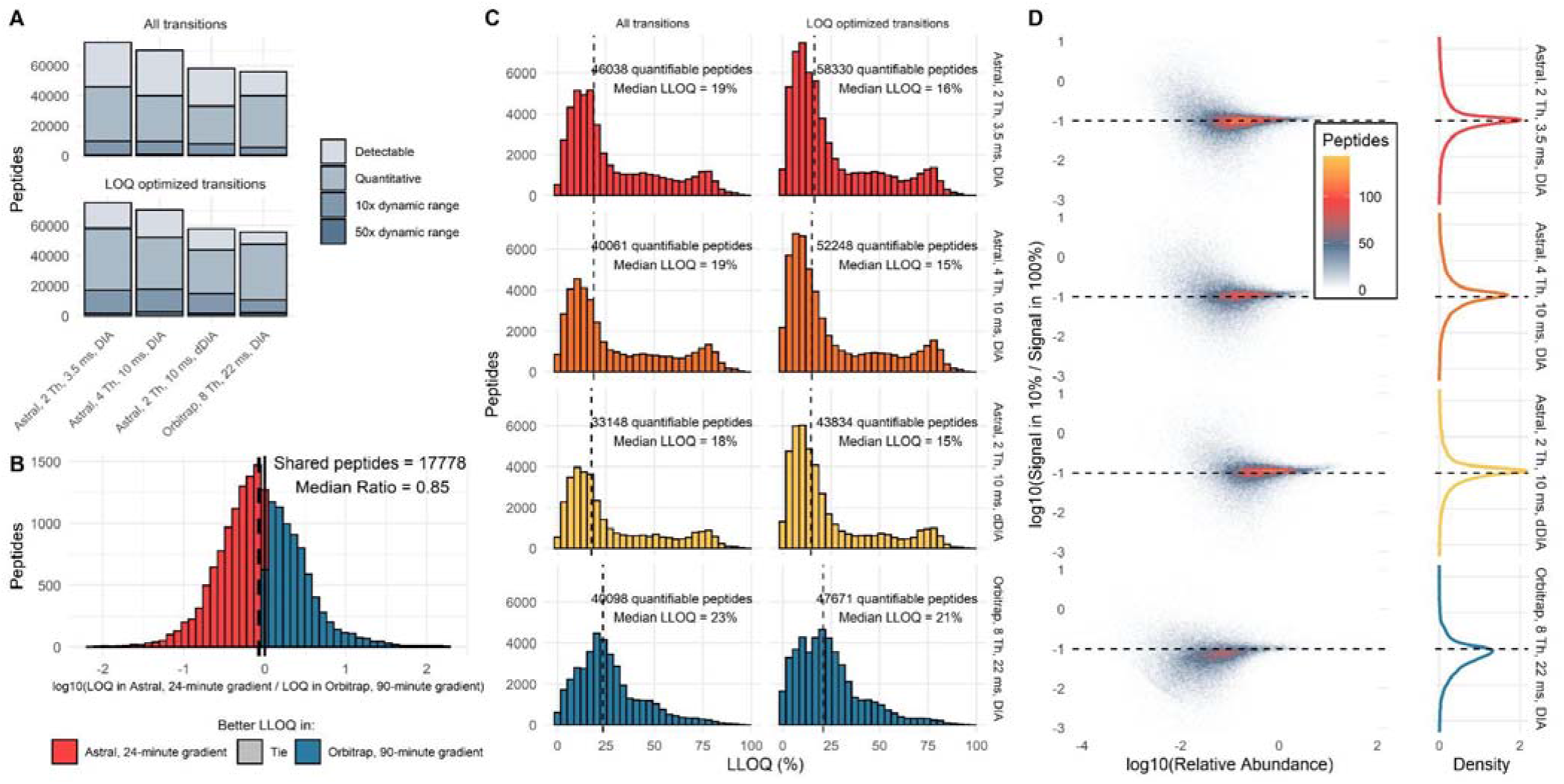
Evaluation of the quantitative performance of the Orbitrap Astral MS. Summary of peptides detected in a matrix-matched calibration curve of HeLa into SILAC labeled HeLa, with 1 µg of total protein load (A). Results of A are summarized in Supplemental Table 2. Peptides are considered detectable if they were detected at 1% FDR, quantitative if they could be assigned a lower limit of quantification less than 100%, quantitative over a 10x dynamic range if that LLOQ was less than 10%, and quantitative over a 50x dynamic range if the LLOQ was less than 2%. Transitions were refined in Skyline to optimize LLOQ and metrics were recalculated with this refined transition set. Pairwise comparison of LLOQs for peptides quantified in the Astral and the Orbitrap, black dashed line represents the median LLOQ (B). Histogram of LLOQs for all quantifiable peptides before and after transition refinement, black dashed lines represent the median LLOQ (C). Signal ratio between 2 points on the dilution curve (100% and 10%) with expected ratio shown as dashed black line, signal from the blank was subtracted from each signal to account for carryover and non-zero background (D). Orbitrap data were acquired with a 90-minute gradient, compared to a 24-minute gradient for the Astral data.

While quantification generally is better in the Astral, the Orbitrap generally performed better across a 100:1 dilution range (**Figure S9**). This trend is consistent with the 24-minute gradient and shorter mass range, where the Orbitrap quantification is slightly better, although fewer peptides are quantified (**Figure S10**). Similarly, the Astral quantifies significantly more peptides (17,454 in Astral with 3.5 ms maximum injection time, 10,686 in Orbitrap) across a 10x linear dynamic range compared to the 90-minute gradient in the Orbitrap, but fewer peptides across a 50x dynamic range (1598 in Astral with 3.5 ms injection time, 1948 in Orbitrap). This relative improvement of Orbitrap performance at lower dilution points may be due to the longer injection times allowing for deeper coverage as well as the higher intra-spectrum dynamic range of the Orbitrap analyzer which is critical for quantification in cases where the peptide of interest is a minor species in a chimeric spectrum. The injection time theory is supported by the fact that quantification in the Astral improved by increasing injection time (2563 peptides have 50x linear dynamic range with 10 ms maximum injection time). In both of these comparisons, the performance of the Orbitrap Astral MS was handicapped by either a shorter gradient or a wider mass range. Even the largest maximum injection time assessed (10 ms) was shorter than the Orbitrap (23 ms). Regardless, the Astral is comparable if not better to the Orbitrap in every method assessed here (**Supplemental Table 2**).

Although the Orbitrap Astral MS has the ability to acquire at rates up to 200 Hz, we predicted that it may be better to use a slower acquisition rate and to increase maximum injection times to increase sensitivity. We found that acquiring spectra at 187 Hz (3.5 ms maximum injection time) and 2 Th isolation windows results in a large number of peptide and protein detections (**Figure 1D**), but these detections tend to have higher CVs and LLOQs than measurements acquired with longer injection times (**Figures 2&3**). However, increasing the maximum injection time to 10 ms required doubling the isolation window to maintain a reasonable cycle time, so we also used a dynamic DIA method which allowed for a 10 ms injection time and 2 Th isolation windows across a variable 300 Th mass range (**Figure 1B-C**). Using dynamic DIA decreases the number of peptide and protein identifications, but results in better technical precision (**Figure 2**) and improved limits of quantification (**Figure 3 and Figure S11**). The longer injection times and narrower isolation windows allowed by dynamic DIA improve quantitative accuracy more as the input gets lower, suggesting that this type of method would be particularly useful in cases where high sensitivity is a priority (**Figure S9**).

We acquired DIA data in the Astral analyzer with a 24-minute gradient at 750 nL/minute as this allows for relatively high throughput. Because the Orbitrap acquires spectra at a much slower rate due to longer detection times and requisite injection times, the dataset we are comparing against was acquired with a 90-minute gradient at 300 nL/minute. There are other differences in the acquisition parameters for the two datasets, such as different columns and a different precursor mass range (380-980 Th in the Astral, 400-1000 Th in the Orbitrap). Therefore, we want to emphasize that this is not a direct comparison, but instead the longer gradient and slower flow rate should put the Orbitrap at a significant advantage. Still, the Astral outperforms the Orbitrap by nearly every quantitative metric while nearly quadrupling the throughput. The Orbitrap has long been the gold-standard in the field for high resolution accurate mass based quantification, and so the improvements in sensitivity and throughput in the Astral analyzer represent a step forward for the field as a whole.

It is notable that the Orbitrap data came from an older dataset acquired on an Orbitrap Fusion Lumos mass spectrometer, which was released 9-years prior to the release of the Orbitrap Astral mass spectrometer. In that time, there have been multiple new instrument releases. Therefore, the Orbitrap dataset described here does not capture peak performance. However, the improvements in performance on modern Orbitrap instruments would likely result in a small relative increase in Orbitrap performance. Guzman et al. found that the Orbitrap Astral MS identified more than twice and many proteins and 3.5x as many precursors as the modern Orbitrap Exploris 480 MS.^27^ Therefore, we would not expect the results to change substantially if a dataset from a more recently released Orbitrap instrument were used for this comparison. Additionally, the MS1 resolution was higher in the Astral analyzer dataset than in the Orbitrap dataset. The MS1 signal is not used for quantitation and therefore likely has little impact on the overall performance of the two instrument platforms.

One limitation of the data reported here is that it was acquired on a prototype instrument running a Beta version of instrument control software. At the time the data were acquired, calibration for the detector gain was undergoing further optimization, and the AGC target of 500% (50,000 charges) may not have been optimal. On the Astral, the major impact of space charging is localized to a given peak. If any single peak contains more than 1,000 ions, the ions begin to repel each other and slow down slightly. As a result, the peak resolution decreases and the mass shifts to a slightly higher value (**Figure S12**). This shift can cause the calibration curve to plateau if it pushes the mass outside of the integration boundary. In extreme cases, the detector signal itself may saturate, but this was not observed with the high dynamic range detector used in the Orbitrap Astral MS.^12^ Overall, we believe that the effects of space charging were relatively minimal here, but could be corrected further by decreasing the AGC target, although this may have a negative impact on LLOQ for some low abundance species that fall in the same isolation window as a more abundant species.

Peptide identification software tools are used to assess whether there is evidence for a specific peptide in a dataset. The false discovery rate is often assessed using either an analytical or empirical null distribution that is determined by the search tool. These tools have gotten very sensitive, often making use of artificial intelligence for the prediction of retention time and fragment ion intensities. However, some of the peptides that the tools assign as being present at a given retention time don’t meet the traditional definition of detectability. By definition a signal is detectable if the measured signal, in this case a background subtracted peak area, is statistically different from a blank. Often, to avoid making multiple measurements, many use a heuristic requiring the signal to be >3x the standard deviation of the background outside of the peak to be detectable. Using this heuristic is further complicated by the fact that different instruments make use of different ion detection and signal processing strategies and have vastly different background signal, making the assessment of whether a signal is detectable near impossible to compare between platforms.^28^ Because of the differences in sensitivity, ion detection principles, and signal processing of the Orbitrap and Astral mass analyzers we used a matched-matrix calibration curve strategy to assess both the LOD and LLOQ for most peptides in a HeLa digest.^14^ We ultimately chose to use the LLOQ because 1) establishing quantitative differences in protein levels is the primary goal of most bottom up proteomics experiments, 2) is a more experimentally relevant measurement than LOD, 3) it would highlight the quantitative performance of each analyzer.

One striking difference between these two high resolution-accurate mass analyzers is the quantity of data produced by the Astral analyzer, as it can acquire spectra much faster than the Orbitrap and each spectrum contains more information, i.e. more centroids per spectrum (**Figure S13**). While we have demonstrated that this extra data contains quantitative information on more peptides, there is also a lot of additional chemical noise that can complicate analysis. Although an in-depth analysis of the extent of this problem is outside the scope of this study, it was evident when visualizing detections from multiple search engines that a significant portion (upwards of 10% in many cases) of the identifications are matching to background signals with no real peak (**Figure S14**). In the majority of these cases, there was extensive peptide backbone coverage generated by low intensity signals, but these signals show little to no correlation across time, meaning that at best they are not providing quantitative information and at worst, they are false matches. We propose that these false identifications are a computational problem that is not unique to the data produced by the Astral analyzer and can be addressed by adapting existing database search strategies.

Notably, 64% of mass spectral peaks in the Astral consist of five or fewer ions (**Figure S15**), although these low intensity noise signals account for only 6.5% of the total ion current. These extra signals in the Astral can interfere with searches if denoising approaches are not used, and tools that were originally designed for Orbitrap data should be carefully evaluated before being applied to data that is not similarly pre-processed. In this experiment we can largely filter out noise by filtering for peptides that are quantitative, meaning they can be assigned a lower limit of quantification where their change in signal is proportional to the change in input. However, there will be many cases where calibration curves are not readily available for a given sample and in many cases a simpler method would be preferable. We propose looking for peptides that have at least 3 co-eluting transitions, a strategy that has been used before for DIA quality control.^22^ In this case, over 76% of quantifiable peptides have at least 3 high quality transitions compared to just 40% of non-quantifiable peptides, suggesting that this type of quality filter may be a good predictor of quantitative behavior. Many of the non-quantitative peptides with high quality chromatographic peaks are contaminants or mis-assigned SILAC labeled peptides that do not respond as expected in the dilution experiment.

While most analyses do not include this type of filtering, we believe that it is critical in reducing noise measurements, and hope that such quality filters become commonplace in the future. We propose that this issue is especially problematic in mass analyzers that are capable of single ion detection, as the spectra are much more complex (**Figure S1C and Figure S2C**) and low-quality identifications are common. In fact, the relative increase in low quality detections (**Figure S14**) could partially explain the high numbers that have been reported recently from time-of-flight instruments. Unlike other popular methods,^29, 30^ the isolation windows used on the Astral analyzer are narrow (2-4 Th), which reduces search space and spectral complexity therefore reducing the chemical noise peaks and the frequency of double-counting peptide features. Although an added ion mobility separation decreases spectral complexity in other methods,^31^ most DIA-PASEF search strategies do not utilize ion mobility to decrease search complexity in the way that narrowing the quadrupole isolation window is able to do. Ultimately, the raw number of peptide and protein identifications alone may be a weak indicator of instrument performance, and should be met with some degree of skepticism until quantitative performance can be validated. The Astral analyzer has higher resolution across the full mass range than the Orbitrap analyzer as operated in these experiments (80k vs 15k) and much higher resolving power than most commercially available TOF instruments. The high resolving power and mass accuracy could help decrease the impact of extraneous signals as a smaller mass tolerance can be used for database searching and peak integration.

### Exploring the plasma proteome

To assess the potential of the Orbitrap Astral MS to generate new biological insights, we tested its performance on plasma, a matrix that is both common and analytically challenging.^2^ While fractionation and enrichment strategies have improved coverage of the plasma proteome, with nanoparticle fractionation detecting 1500-2000 proteins across 5 runs^32^ and other methods yielding up to 2700 proteins in a single run,^33^ the plasma proteome still remains a formidable analytical challenge. The two main ways to address the dynamic range problem are to increase the dynamic range of the mass spectrometer itself or to use an enrichment or depletion strategy that will reduce the dynamic range of the samples we are measuring. Here, we apply both strategies by using the highly sensitive Orbitrap Astral MS to measure plasma proteins from an extracellular vesicle enriched sample.

The sensitivity and quantitative dynamic range of the Astral analyzer is ideally suited for complex biofluids such as plasma. The dynamic range of plasma is especially challenging for mass spectrometers as electron multiplier detectors may only have a linear dynamic range of 3-4 orders of magnitude, which is much smaller than the dynamic range of plasma. Unlike TOFs, the Orbitrap Astral MS uses automatic gain control of ion injection times, which can expand the dynamic range by two or more orders of magnitude. Therefore, we expected the Astral analyzer to perform especially well on plasma and quantify proteins across a wide dynamic range.

To reduce dynamic range, we used a simple magnetic bead-based protocol that is capable of enriching for extracellular vesicles (EV) from plasma. Because most of the abundant plasma proteins are not associated with or within vesicles they are depleted during the EV enrichment process.^16^ The EVs are enriched using a combination of the MagResyn hyper-porous polymer matrix that functions as a molecular “net” for membrane bound particles. The polymer matrix of the beads contains a quaternary ammonium surface chemistry resulting in a cationic charge (MagResyn SAX bead). The porous polymer allows vesicles and biomolecules to intercalate within the volume of the beads and positive charge provides a way to distinguish vesicles from lipoprotein particles based on the charge.^34^ This protocol results in the enrichment of known EV markers by ∼20-fold and the depletion of common plasma proteins by ∼95%. Using this protocol, Wu et al. were able to improve protein detections from 1,088 in total plasma (using a library generated from plasma EVs) to 4,163 in enriched plasma.^16^

To evaluate plasma proteome coverage in the Astral-based instrument we assessed the quantitative performance of multiple acquisition methods with a short (30-minute) and long (60-minute) gradient in both EV enriched and total plasma. We used multiple isolation window sizes and injection times with each gradient to assess the merits of these parameters. For the 30-minute gradient, we operated the instrument at 187 Hz (3.5 ms maximum injection time, 2 Th isolation windows), 90 Hz (10 ms maximum injection time, 4 Th isolation windows), and 50 Hz (20 ms maximum injection time, 4 Th dynamic DIA windows). For the 60-minute gradient, we tested two methods both operating at 60 Hz (15 ms maximum injection time), one using 4 Th static isolation windows and one using 2 Th dynamic isolation windows.

Unsurprisingly, we found that the longer gradient produced more peptide and protein level identifications than the shorter gradient (**Figure 4A**). The 60-minute gradient with 4 Th isolation windows and a 15 ms maximum injection time yielded the most peptide and protein-level detections, with 44,668 peptides from 5,163 proteins (**Figure 4C)**. Using a 30-minute gradient with 4 Th isolation windows and a 10 ms maximum injection time, we detected 37,534 peptides from 4,704 proteins. Generally, slowing down acquisition to allow for longer injection times led to better technical precision, although cutting the mass range for dynamic DIA reduced the number of peptide and protein detections. All methods assessed here detected over 4200 proteins with high technical precision, suggesting that the Astral is well-suited for these types of measurements (**Figure 4B**). Further, the Astral is able to detect enrichment and depletion of certain marker proteins that span a wide dynamic range (**Figure S16**) relative to total plasma (**Figure 4D**).^16^ Using the same enrichment protocol, Wu et al. detected 37,942 peptides from 4,163 proteins on a Thermo Scientific Orbitrap Eclipse MS with a 110-minute gradient.^16^ The Orbitrap Astral MS is able to detect more proteins and a similar number of peptides in a 30-minute gradient, highlighting that one benefit of the Astral analyzer’s increased speed and sensitivity is the ability to increase throughput while maintaining or even increasing coverage.

**Figure 4.**
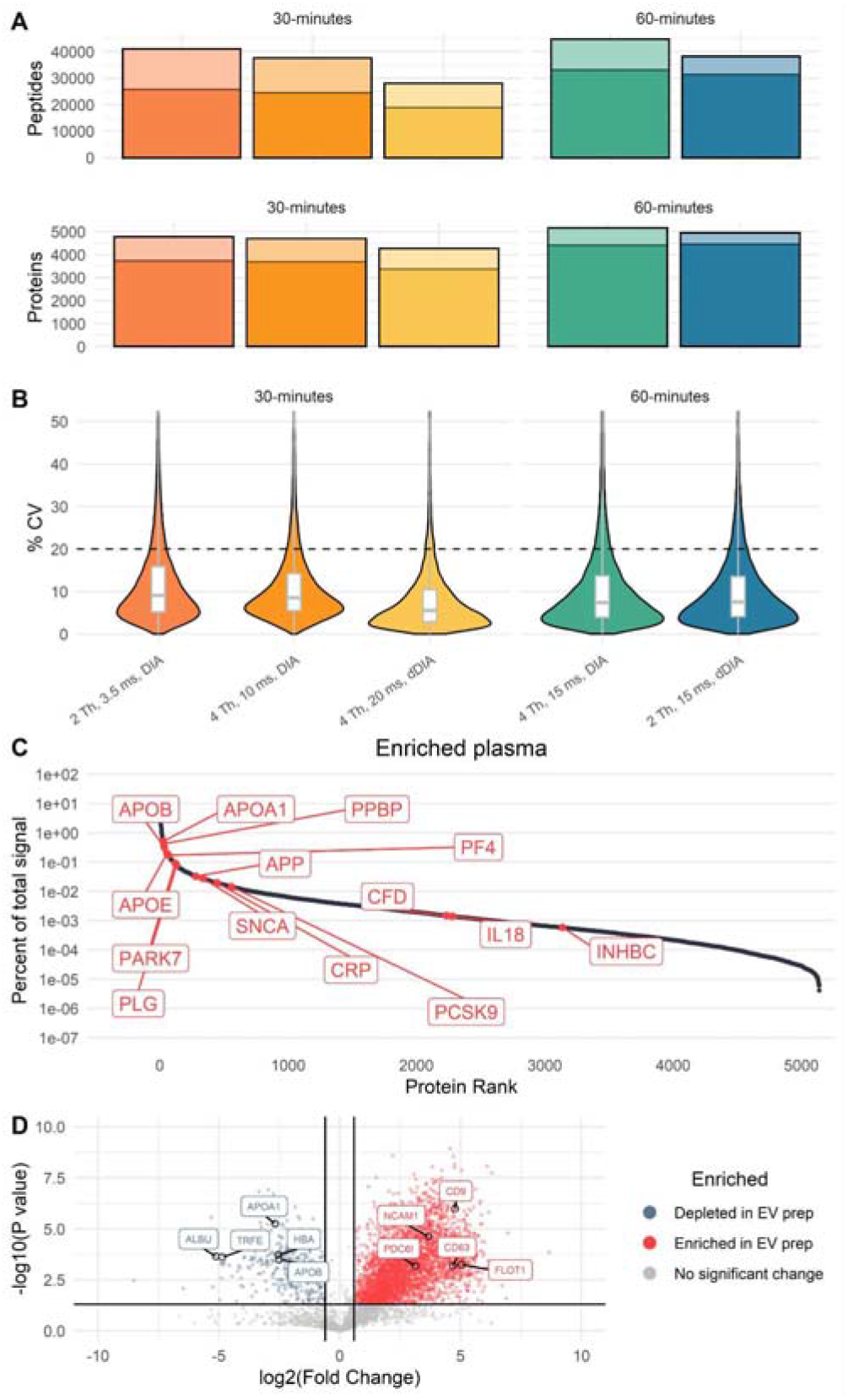
Plasma protein quantification. Peptide and protein-level detections from plasma samples (A). Lighter bars represent peptides/ proteins that were detected at 1% FDR but did not have a chromatographic peak with at least 3 co-eluting transitions. Summary of all proteins measured with selected biologically relevant proteins highlighted (B). Coefficient of variation of all peptides detected in EV-enriched plasma with each method (C). The black dashed line is at 20% CV; in all cases more than 83% of all peptides had CVs less than 20%. Volcano plot of proteins enriched and depleted in the EV-enrichment method relative to a total plasma preparation, results are shown for the 60-minute, 4 Th, 15 ms DIA method, and are representative for the results from all five methods (D). Vertical black lines indicate a fold change of +/-1.5, horizontal black line indicates a significance threshold of 0.95.

While technical precision and concordance with known biology suggest that the measurements produced by the Orbitrap Astral MS are meaningful, it was evident that a certain proportion of these detections can be attributed to background signals and chemical noise (**Figure S14**). Thus, to determine the proportion of high-quality peptides and proteins, we filtered the results for peptides with at least three co-eluting transitions. After applying this filter, the number of peptides and proteins decreased only slightly to a maximum of 31,373 peptides from 4,453 proteins with the dynamic DIA method (**Figure 4A**). One result of filtering for peptides with multiple co-varying transitions is an elimination of peptides with chemical interference leading to an improved coefficient of variation (**Figure S17**). Here, increasing the injection time improves the ion statistics which increases both the sensitivity and the precision. Likewise, narrowing the isolation windows improves the measurement selectivity and reduces chemical interference. Both the increased injection time and improved selectivity leads to improved detection limits (**Figure 4A**). Even with a relatively strict quality filter, the plasma proteome coverage presented here is unprecedented.

## Conclusions

Overall, this initial quantitative characterization suggests that the Orbitrap Astral mass spectrometer generates high quality, quantitative data for a large number of peptides. The high acquisition rate, high resolution, sensitivity, and the power of automatic gain control produce previously unattainable dynamic range across the proteome. In our analysis, the Astral analyzer quantified more peptides and proteins than the Orbitrap could in one fourth of the time, meaning that with the Orbitrap Astral mass spectrometer the analytical throughput can be quadrupled without sacrificing quantitative performance. Additionally, the increased speed of the Astral analyzer allows for smaller isolation windows while covering the same mass range, producing spectra that are less complex and computationally easier to search. These advances are critical for maintaining precision and accuracy while increasing throughput of measurements.

Our results also highlight the need for quantitative characterization and careful consideration of an instrument’s strengths when selecting the best instrument for the project. Despite all of the strengths of the Astral, the intra-spectrum dynamic range of the Orbitrap is superior, albeit takes a longer acquisition time to achieve, and it is important to select the appropriate analyzer for a given measurement based on the specific goals and practical limitations. Here, we use calibration curves spanning a wide dynamic range to determine the lower limit of quantification for tens of thousands of peptides in addition to limits of detection and coefficients of variation. These figures of merit provide much more context and meaning than simply evaluating protein-level identifications or even looking at a single three-proteome mixture, as has become the common practice in the field.

Not only is the quantitative performance of the Orbitrap Astral mass spectrometer analytically robust, but we have demonstrated the potential to obtain improved detection of low abundance proteins in the plasma proteome when combined with a novel, yet simple, protocol.^16^ We can now quantify over 5000 plasma proteins in a single one-hour LC-MS/MS run (**Figure 4C**) which was previously unattainable. This depth and quality of coverage is just one application where speed, sensitivity, and dynamic range of the new analyzer enable improved quantitative coverage in a shorter time span.

## Notes

The authors declare the following competing financial interest(s): The MacCoss Lab at the University of Washington has a sponsored research agreement with Thermo Fisher Scientific, the manufacturer of the instrumentation used in this research. However, analytical techniques were selected and performed independent of Thermo Fisher Scientific. M.J.M. is a paid consultant for Thermo Fisher Scientific. E.D.,T.N.A., A.C.P., E.D., J.P., A.P., P.M.R., M.W.S., H.I.S., C.H., A.A.M., D.H., and V.Z. are employees of Thermo Fisher Scientific, the manufacturer of the instrumentation used in this research.

## Acknowledgements

This work was supported in part by National Institutes of Health grants U19 AG065156, R24 GM141156, and P30 AG013280.

---

Supplemental File S1: Supplemental figures referenced in text

Figure S1: Ion statistics for HeLa chromatogram library, 24 minute gradient

Figure S2: Ion statistics for quantitative DIA methods

Figure S3: UpSet plot of peptide detections across methods

Figure S4: UpSet plot of peptide detections across a limited mass range, same LC method

Figure S5: Peptide and protein CVs across limited mass range

Figure S6: Peptides quantified per unit time

Figure S7: Quantification across a 10x dilution using standard protein grouping

Figure S8: Quantification across a shorter mass range

Figure S9: Quantification across 2 orders of magnitude

Figure S10: Quantification across 2 orders of magnitude, shortened mass range

Figure S11: Effects of dynamic DIA on limit of quantification

Figure S12: Mass error vs number of ions

Figure S13: Selected fragmentation spectrum in Orbitrap compared to Astral analyzer

Figure S14: Selected examples of low abundance peptides detected in the Astral data using different software tools

Figure S15: Ions per peak

Figure S16: Plasma dynamic range

Figure S17: Plasma CVs before and after refinement

Supplemental File S2: Supplemental tables referenced in text

Table S1: Summary of LC-MS/MS runs

Table S2: Summary of peptide detections in matrix matched calibration curve

## FOR TOC ONLY

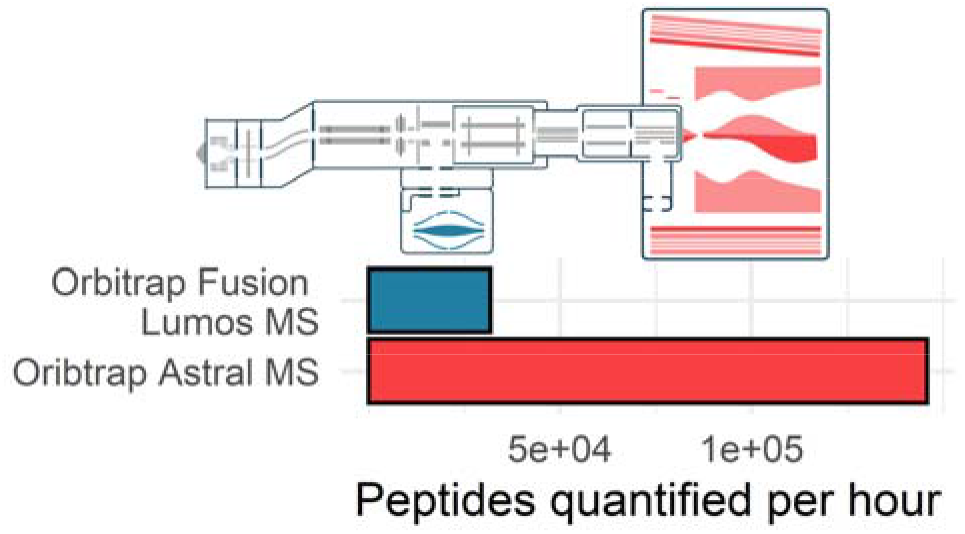

This TOC figure was created by the authors specifically for this manuscript and has not been published elsewhere.

