## Supplemental Figures for "Evaluating the Performance of the Astral Mass Analyzer for Quantitative Proteomics Using Data Independent Acquisition"



Figure S1: Ion statistics for HeLa chromatogram library, 24 minute gradient
Figure S2: Ion statistics for quantitative DIA methods
Figure S3: UpSet plot of peptide detections across methods
Figure S4: UpSet plot of peptide detections across a limited mass range, same LC method
Figure S5: Peptide and protein CVs across limited mass range
Figure S6: Peptides quantified per unit time
Figure S7: Quantification across a 10x dilution using standard protein grouping
Figure S8: quantification across a shorter mass range
Figure S9: Quantification across 2 orders of magnitude
Figure S10: Quantification across 2 orders of magnitude, shortened mass range
Figure S11: Effects of dynamic DIA on limit of quantification
Figure S12: Mass error vs number of ions
Figure S13: Selected fragmentation spectrum in Orbitrap compared to Astral analyzer
Figure S14: Selected examples of low abundance peptides detected in our Astral data using different software tools
Figure S15: Ions per peak
Figure S16: Plasma dynamic range
Figure S17: Plasma CVs before and after refinement

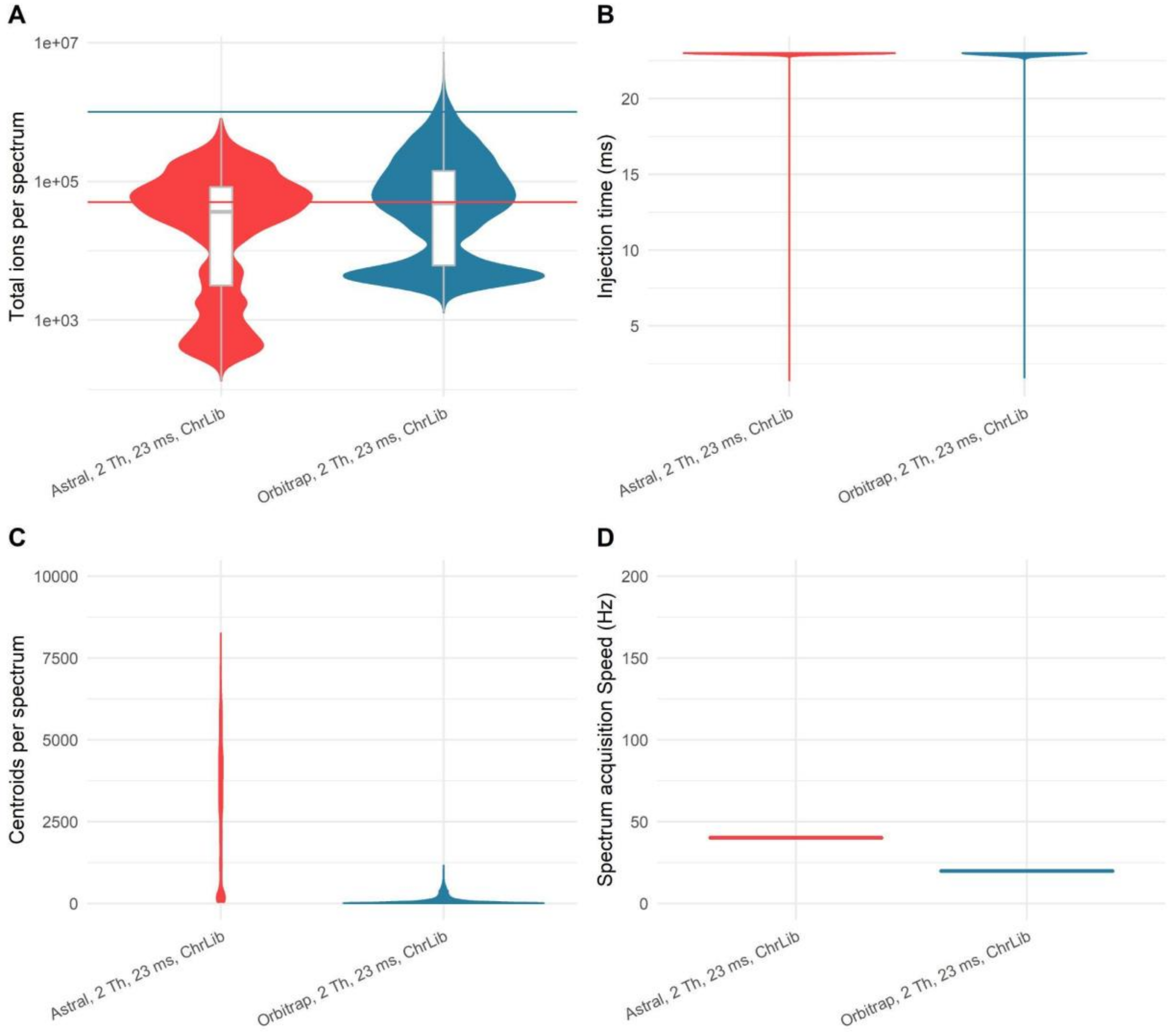


**Figure S1**. *Ion statistics for HeLa chromatogram library, 24 minute gradient*. Comparison of ions per spectrum (A), injection time (B), centroids per spectrum (C), and overall acquisition speed for both Orbitrap and Astral in a 24-minute chromatogram library run.

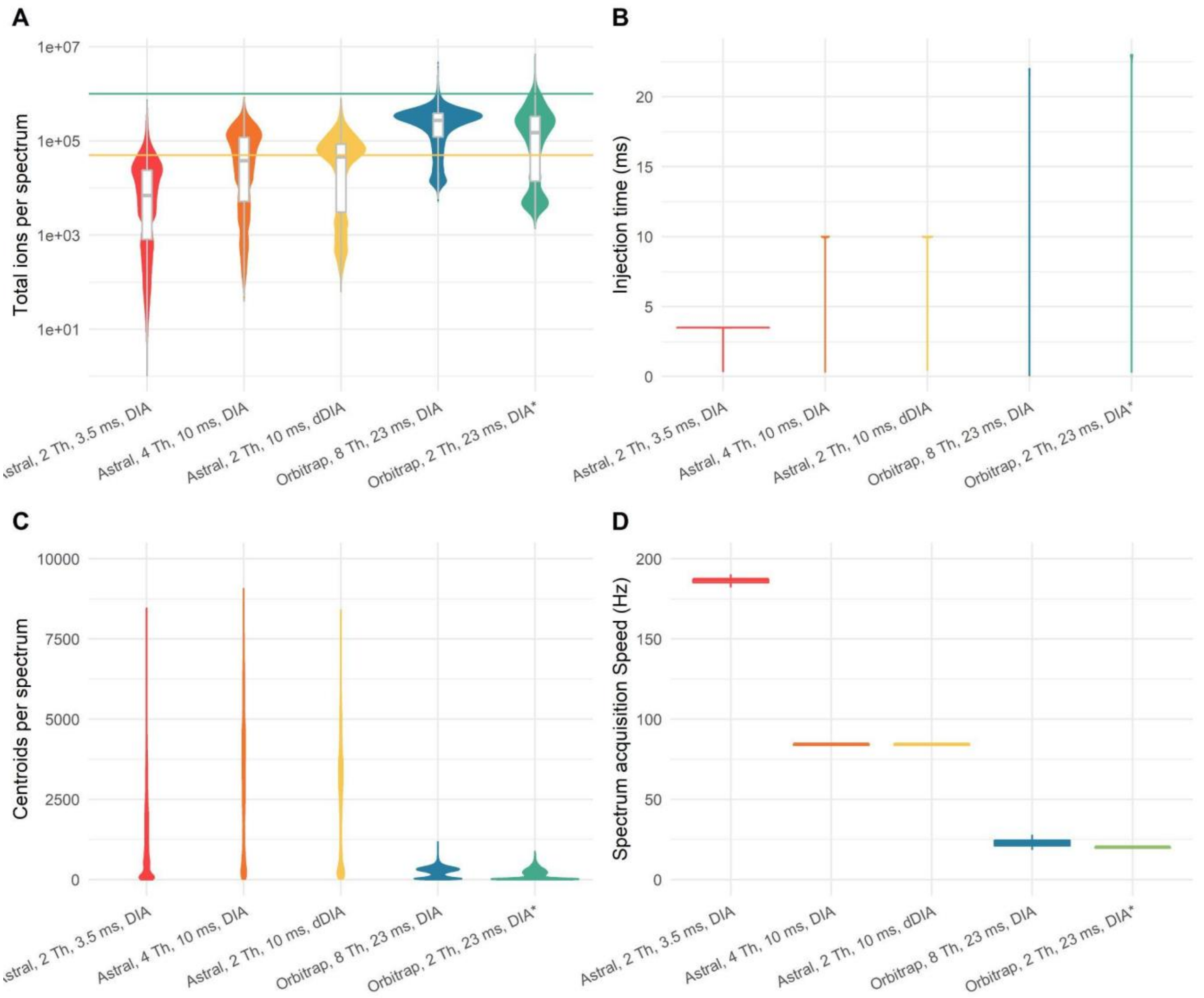


**Figure S2**. *Ion statistics for quantitative DIA methods*. Comparison of ions per spectrum (A), injection time (B), centroids per spectrum (C), and overall acquisition speed for both 3 Astral methods acquired with a 24-minute gradient, one Orbitrap method acquired with a wide mass range and a 90-minute gradient, and one Orbitrap method acquired with a 75 Th mass range and the same, 24-minute gradient used in the Astral methods.

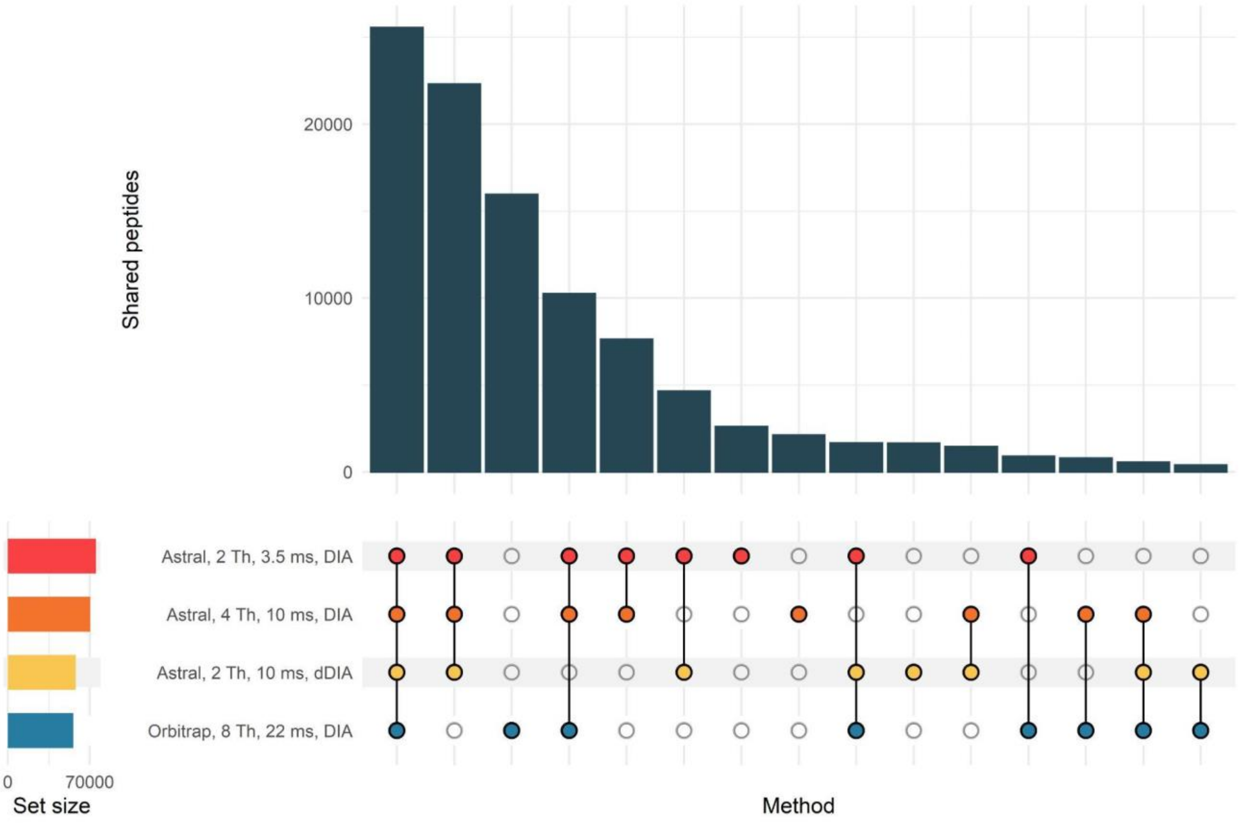


**Figure S3**. *UpSet plot of peptide detections across methods*. Comparison of peptide-level detections in 1 µg of HeLa digest in the Astral with a 24-minute gradient and the Orbitrap with a 90-minute gradient.

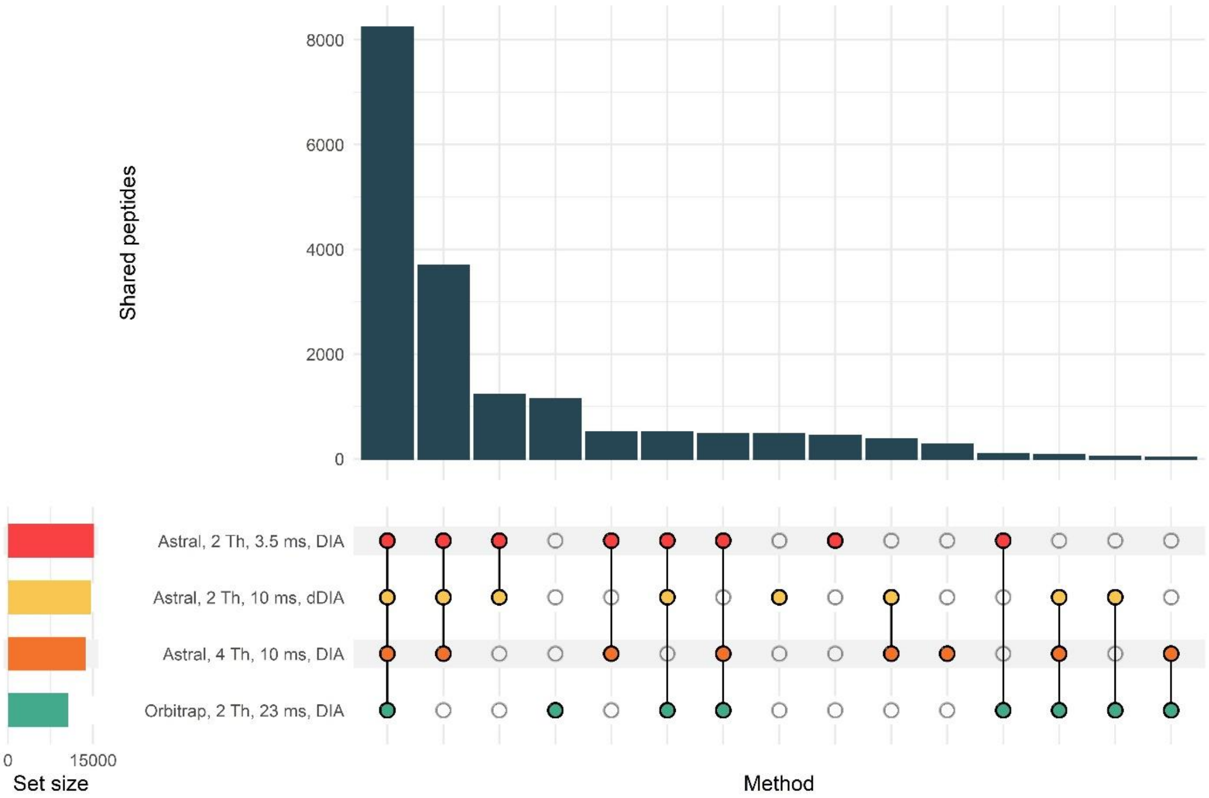


**Figure S4.** *UpSet plot of peptide detections across a limited mass range, same LC method*. Comparison of peptide-level detections in 1 µg of HeLa digest in the Orbitrap and the Astral with a 24-minute gradient. The Orbitrap method acquired DIA spectra across a limited (75 Th) mass range, and the Astral data was originally acquired across a larger mass range but was filtered down for the analysis to peptides in the same 75 Th mass range.

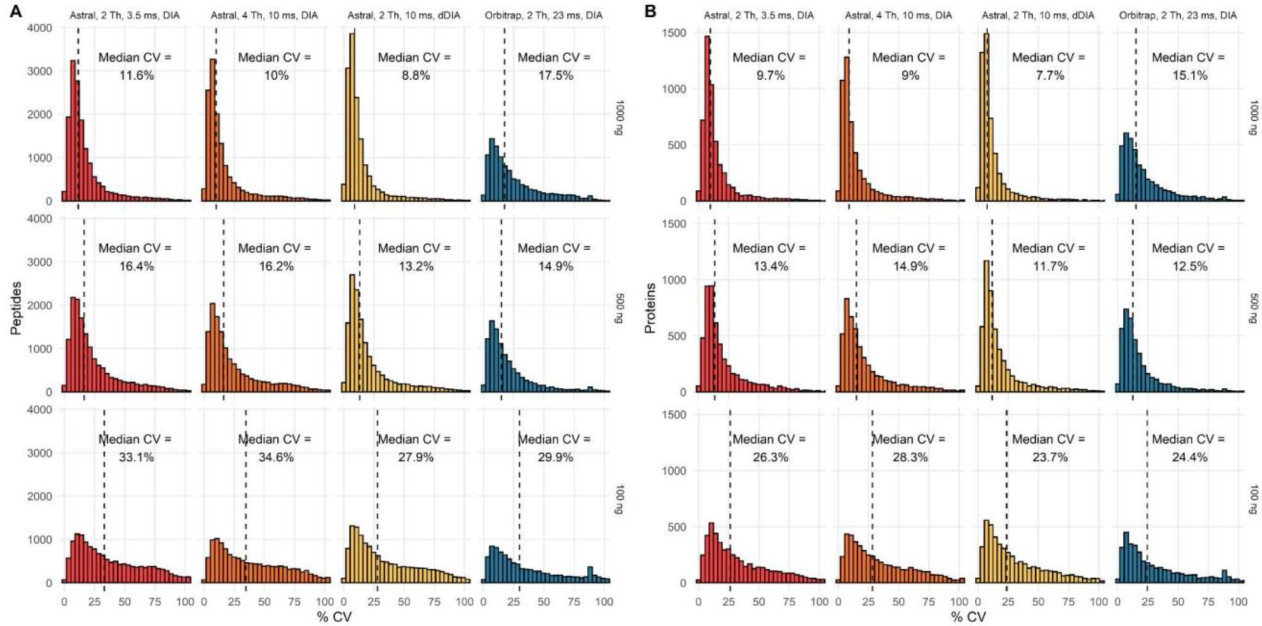


**Figure S5.** *Peptide and protein CVs across limited mass range*. Peptide (A) and protein (B) level coefficients of variation across 4 different acquisition methods. CVs are plotted for the same set of peptides with different amounts of HeLa injected with SILAC HeLa added as a background to keep total protein loading level constant (1000 ng). Dashed horizontal line represents the median CV, which is also indicated on the plot. The Orbitrap method acquired DIA spectra across a limited (75 Th) mass range, and the Astral data was originally acquired across a larger mass range but was filtered down for the analysis to peptides in the same 75 Th mass range.

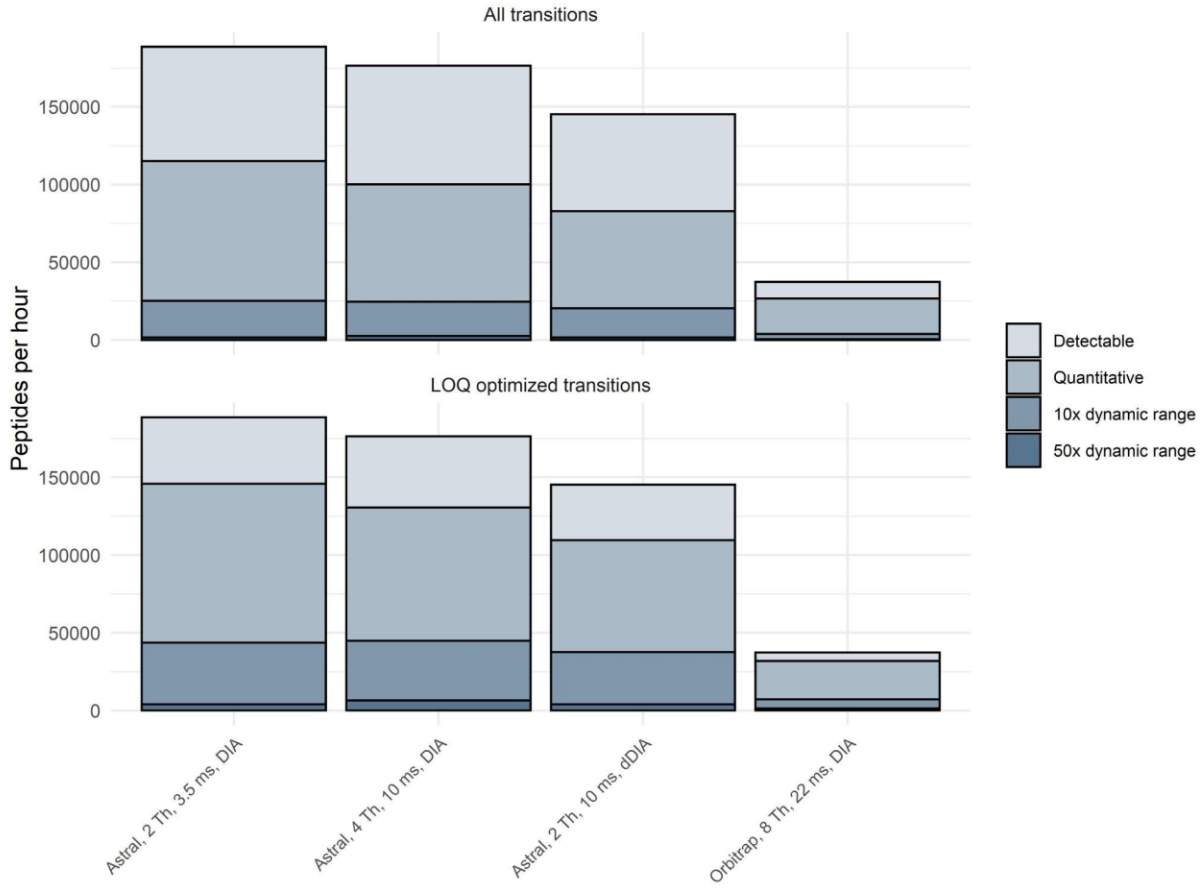


**Figure S6.** *Peptides quantified per unit time.* Plot of peptides quantified per hour. Same data shown in Figure 3A and Table S2, scaled to gradient length.

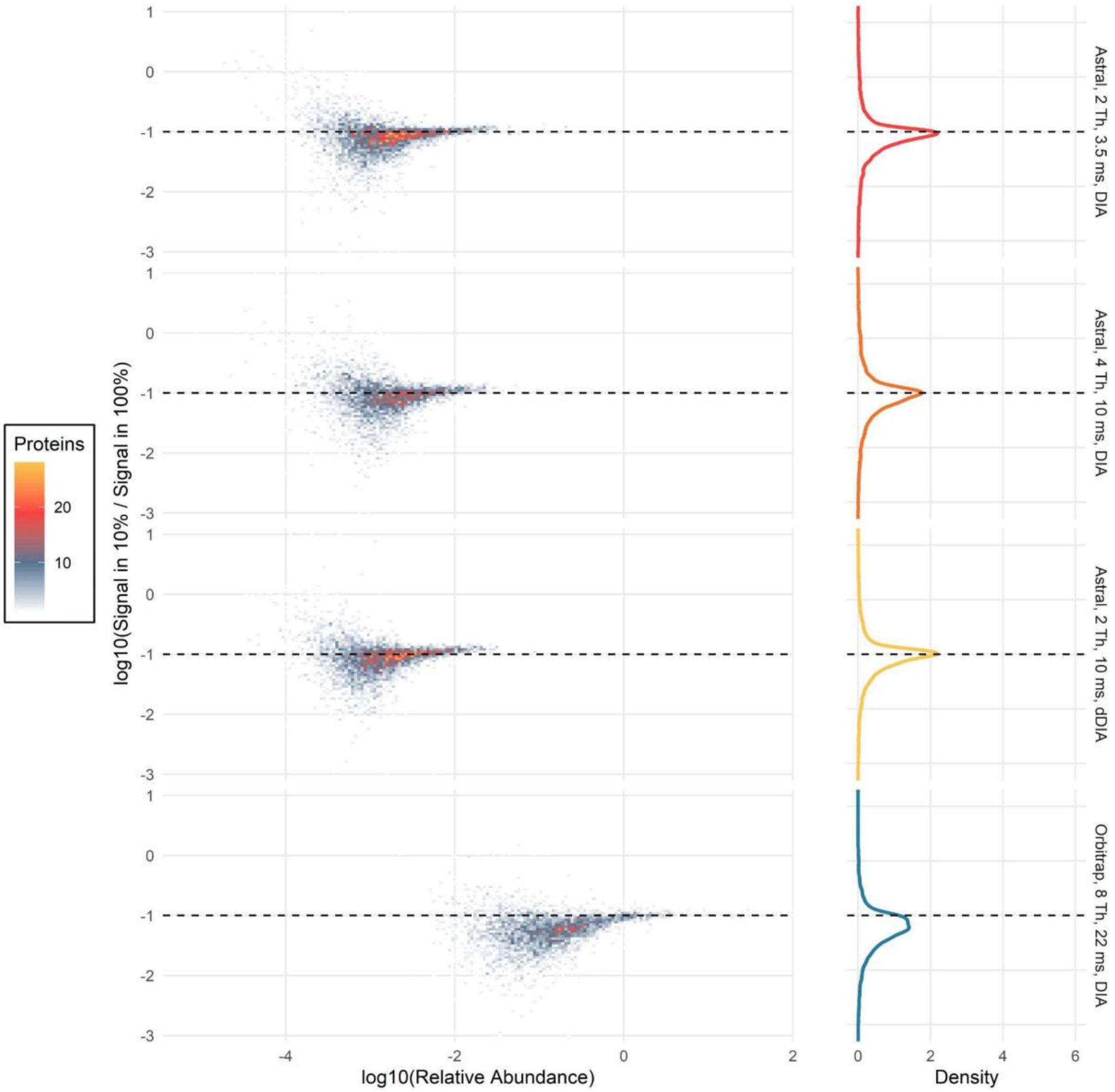


**Figure S7**. *Quantification across a 10x dilution using standard protein grouping*. Density plot of quantitative ratios across a 10x dilution using protein grouping. Astral data were acquired with a 24-minute gradient, compared to Orbitrap which used a 90-minute gradient.

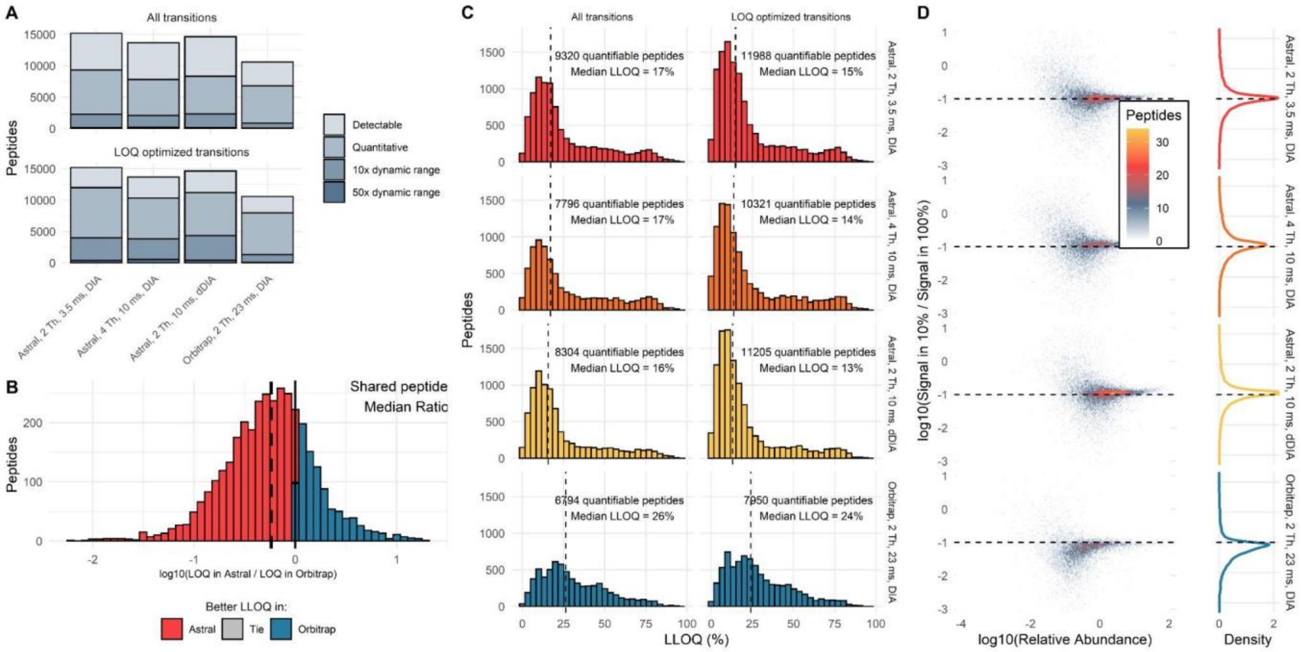


**Figure S8.** *Quantification across a shorter mass range*. Summary of peptides detected in a matrix-matched calibration curve of HeLa into SILAC labeled HeLa, with 1 µg of total protein load (A). Peptides are considered detectable if they were detected at 1% FDR, quantitative if they could be assigned a lower limit of quantification less than 100%, quantitative over a 10x dynamic range if that LLOQ was less than 10%, and quantitative over a 50x dynamic range if the LLOQ was less than 2%. Transitions were refined in Skyline to optimize LLOQ and metrics were recalculated with this refined transition set. Pairwise comparison of LLOQs for peptides quantified in the Astral and the Orbitrap, black dashed line represents the median LLOQ (B). Histogram of LLOQs for all quantifiable peptides before and after transition refinement, gray dashed lines represent the median LLOQ (C). Signal ratio between 2 points on the dilution curve (100% and 10%) with expected ratio shown as dashed black line (D). The Orbitrap method acquired DIA spectra across a limited (75 Th) mass range, and the Astral data was originally acquired across a larger mass range but was filtered down for the analysis to peptides in the same 75 Th mass range.

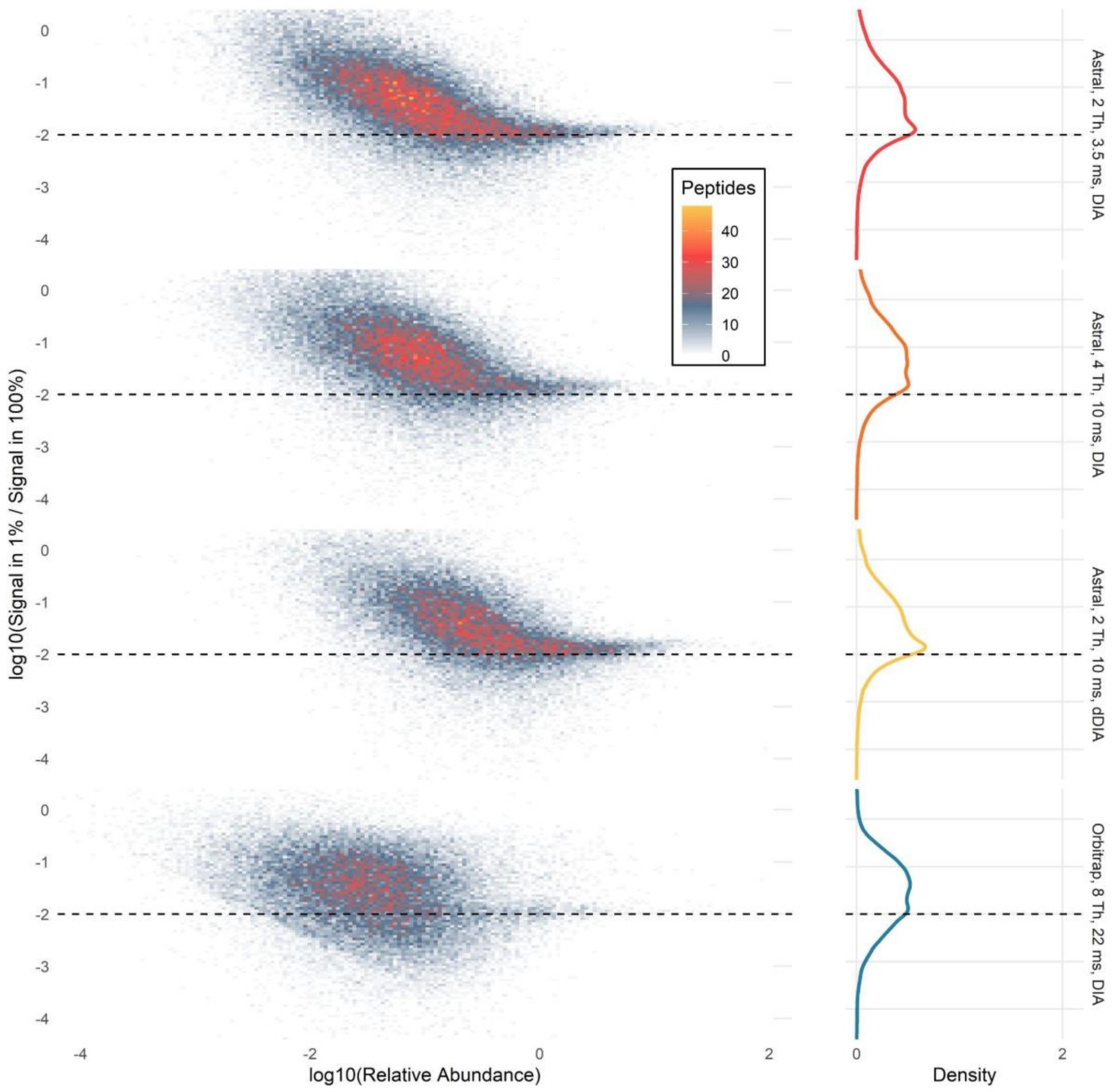


**Figure S9**. *Quantification across 2 orders of magnitude*. Density plot of peptide quantitative ratios across a 100x dilution. Astral data were acquired with a 24-minute gradient, compared to Orbitrap which used a 90-minute gradient.

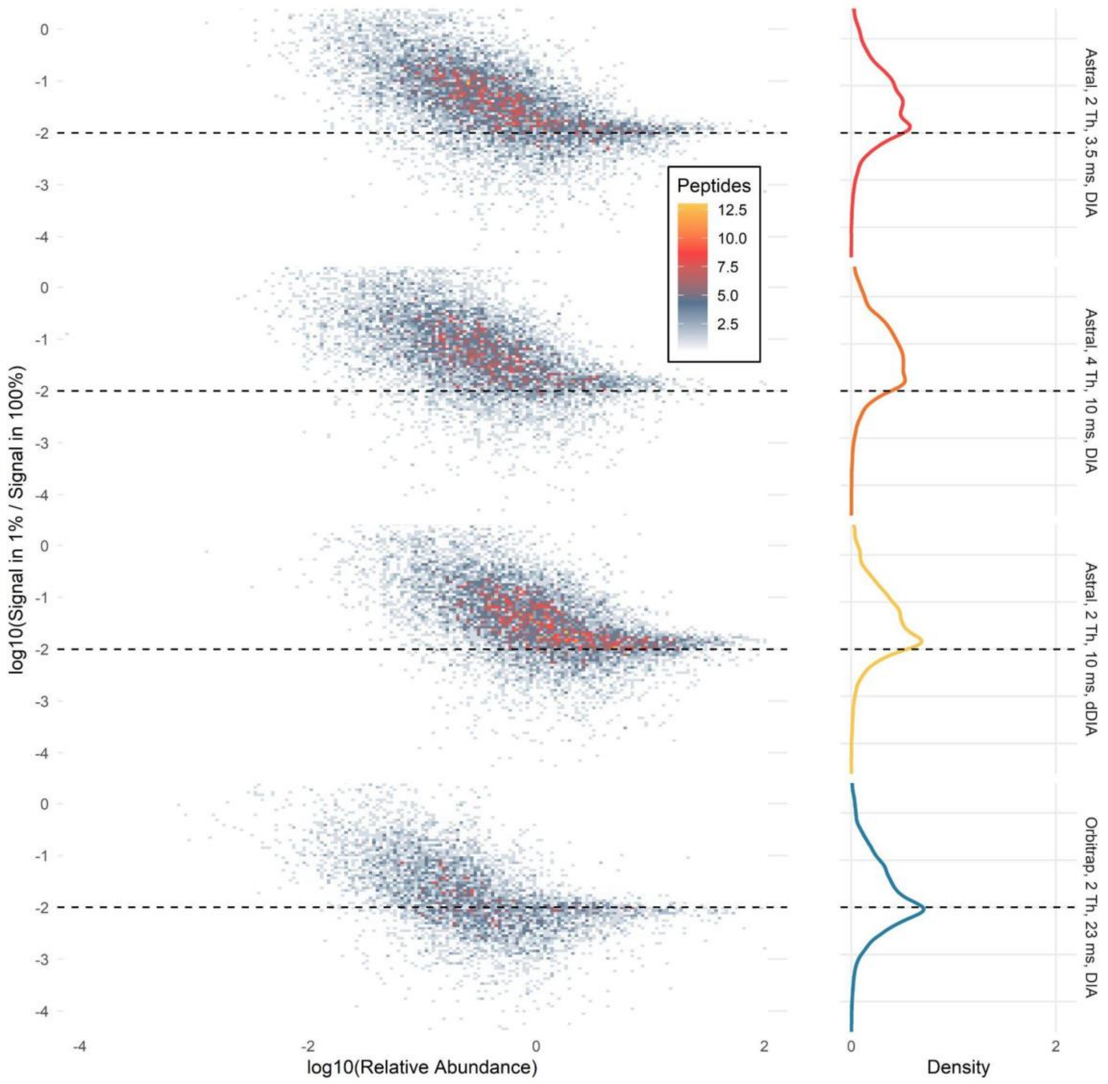


**Figure S10**. *Quantification across 2 orders of magnitude, shortened mass range*. Density plot of peptide quantitative ratios across a 100x dilution. Astral data were acquired with a 24-minute gradient, compared to Orbitrap which used a 90-minute gradient. The Orbitrap method acquired DIA spectra across a limited (75 Th) mass range, and the Astral data was originally acquired across a larger mass range but was filtered down for the analysis to peptides in the same 75 Th mass range.

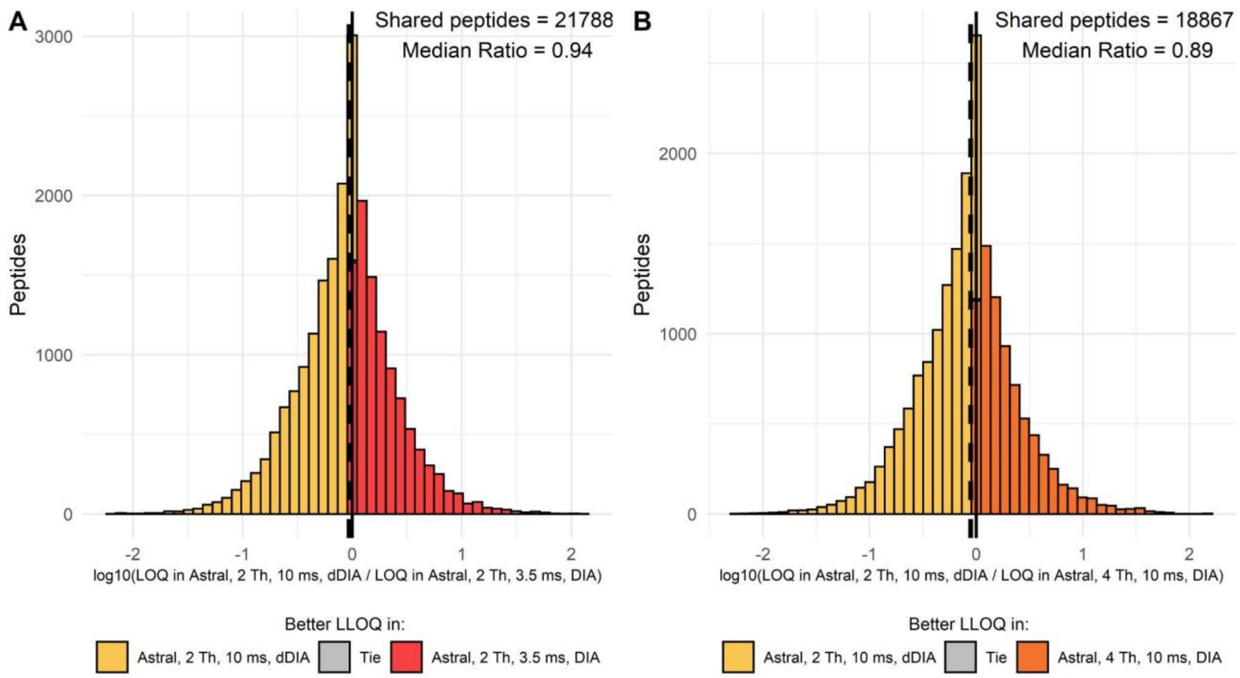


**Figure S11**. *Effects of dynamic DIA on limit of quantification*. Pairwise comparison of lower limits of quantification obtained by peptides quantified in multiple methods by the Astral. Dashed black line represents the median ratio. A log ratio greater than 0 means that the dynamic DIA method was more sensitive than the static acquisition method.

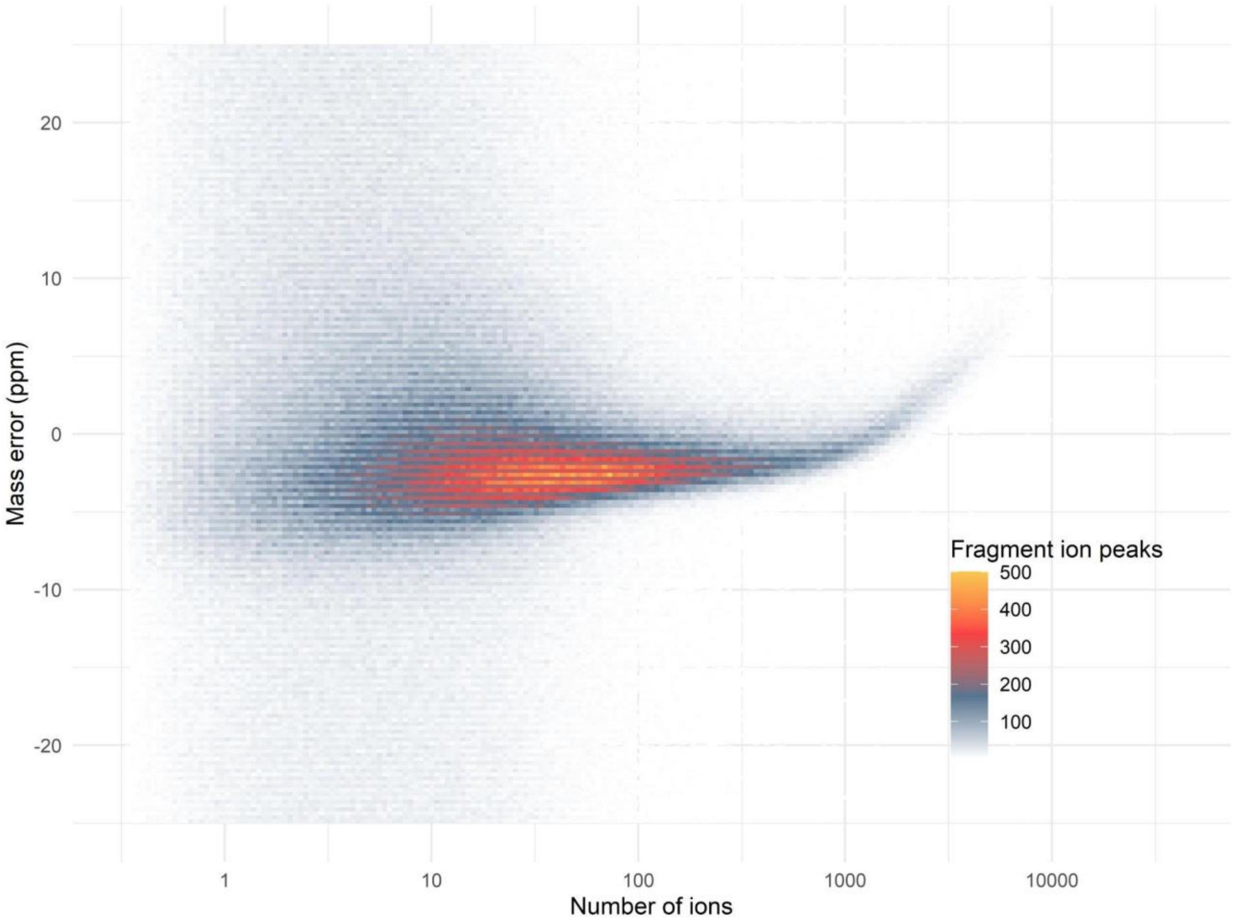


**Figure S12**. *Mass error vs number of ions*. 2-dimensional density plot of fragment mass error vs number of ions. Mass errors are shown for every fragment ion in each point across the peak. A clear inflection point can be seen above 1000 ions which is consistent with localized space charging.

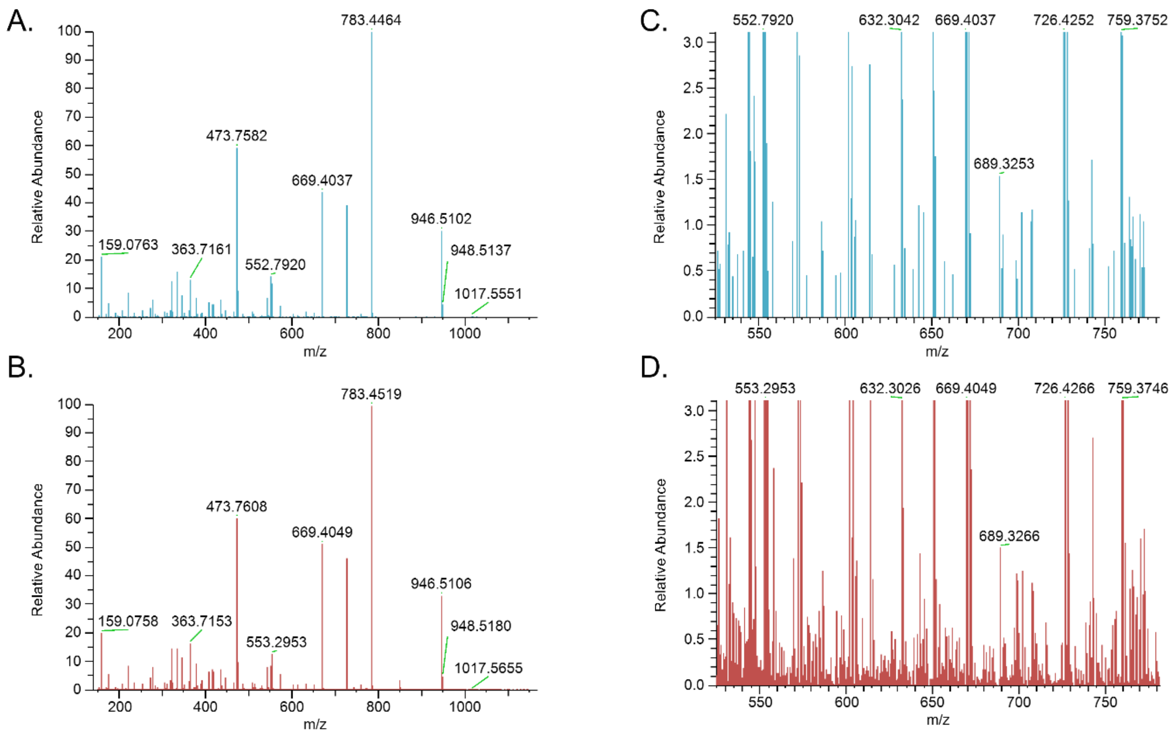


**Figure S13**: *Selected fragmentation spectrum in Orbitrap compared to Astral analyzer*. A spectrum covering the same mass range at the same retention time in the Orbitrap (A, C) and Astral (B, D) analyzers. A and B are full spectra, C and D are zoomed in on a section of low intensity. While the full spectra look similar, there are additional low intensity peaks in the spectrum acquired in the Astral analyzer.

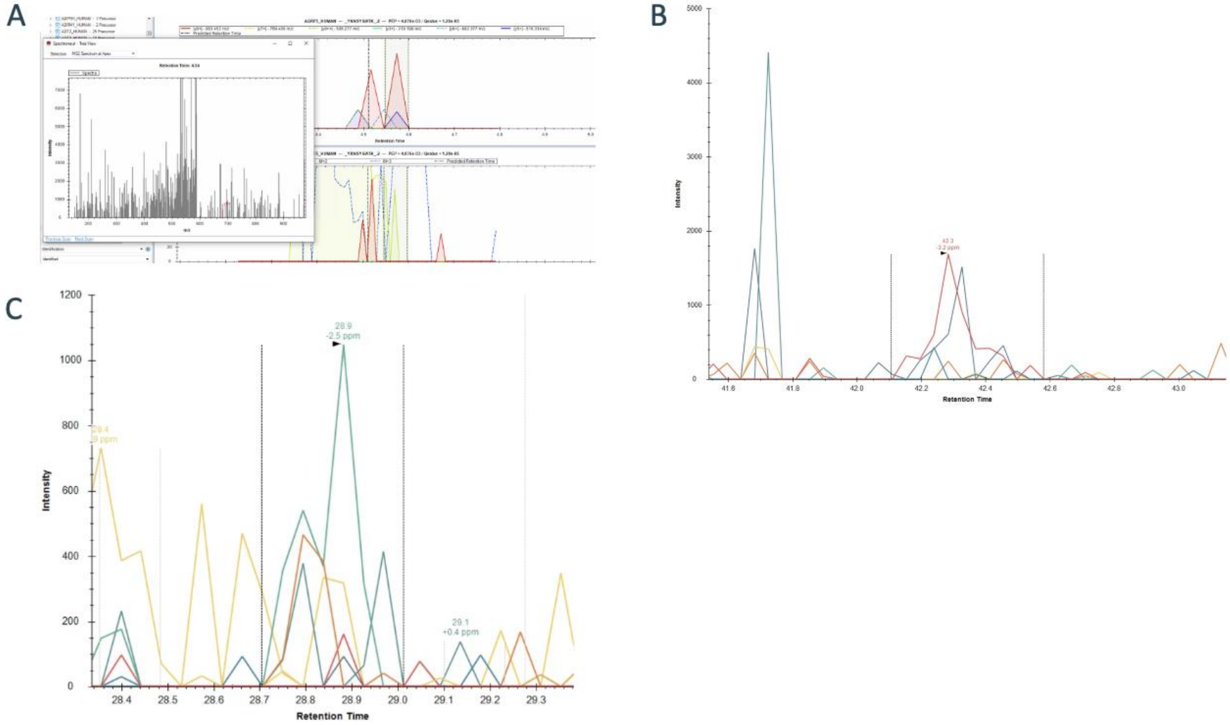


**Figure S14.** *Selected examples of low abundance peptides detected in our Astral data using different software tools*. These are example peptides detected by Spectronaut (A), EncyclopeDIA (B), and DIA-NN (c) respectively with no clear chromatographic peak. Despite being called "identified," these peptides are likely below the limit of detection (LOD). Peptides matching to low abundance signals can account for up to 10% of all detections regardless of the search tool used. Each tool has different scoring algorithms, methods for assessing the FDR, and strategies to account for background. Comparing different instrument platforms that use different ion signal detection and signal processing methods (e.g. the Orbitrap and Astral) is complicated. Differences in the background signal written to the raw datafile make the assessment of S/N difficult between instruments (Wells et al., Application Note 2023). Because of these challenges, we use at matrix-matched calibration curve strategy to compare our Orbitrap and Astral analyzer quantitative data (Pino et al, J. Proteome Res 2020). This strategy uses many samples at known and different relative quantities to assess the LOD and LLOQ.

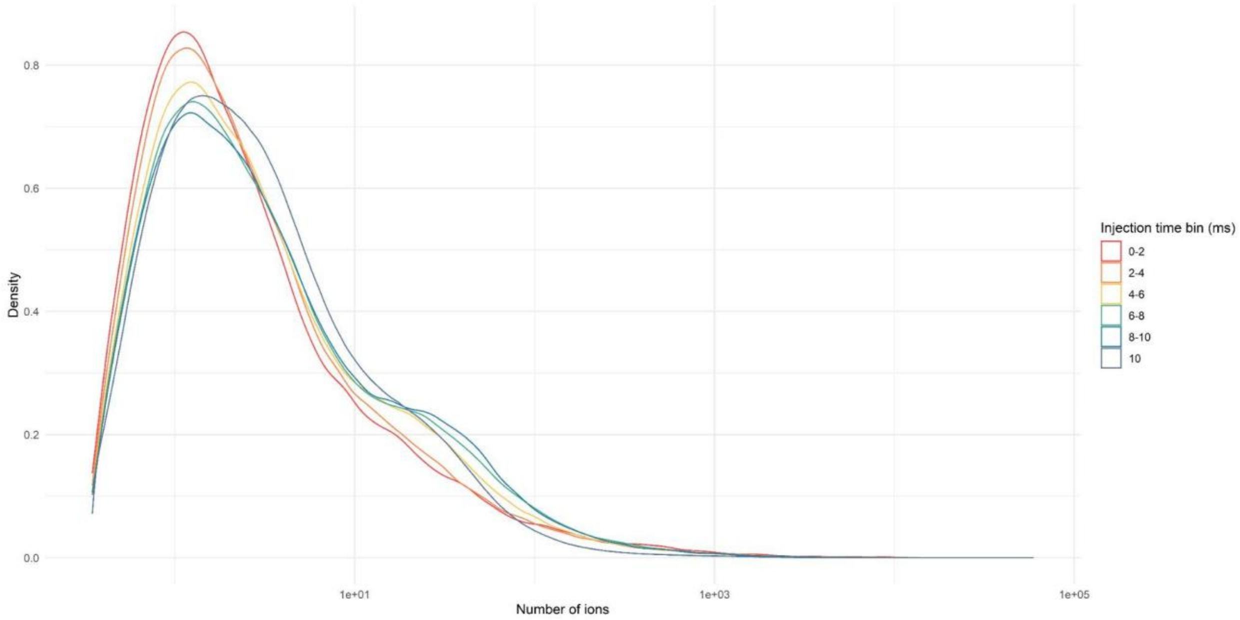


**Figure S15**. *Ions per peak.* Density plot of ions per spectral peak in the Astral. Spectra were binned by injection time.

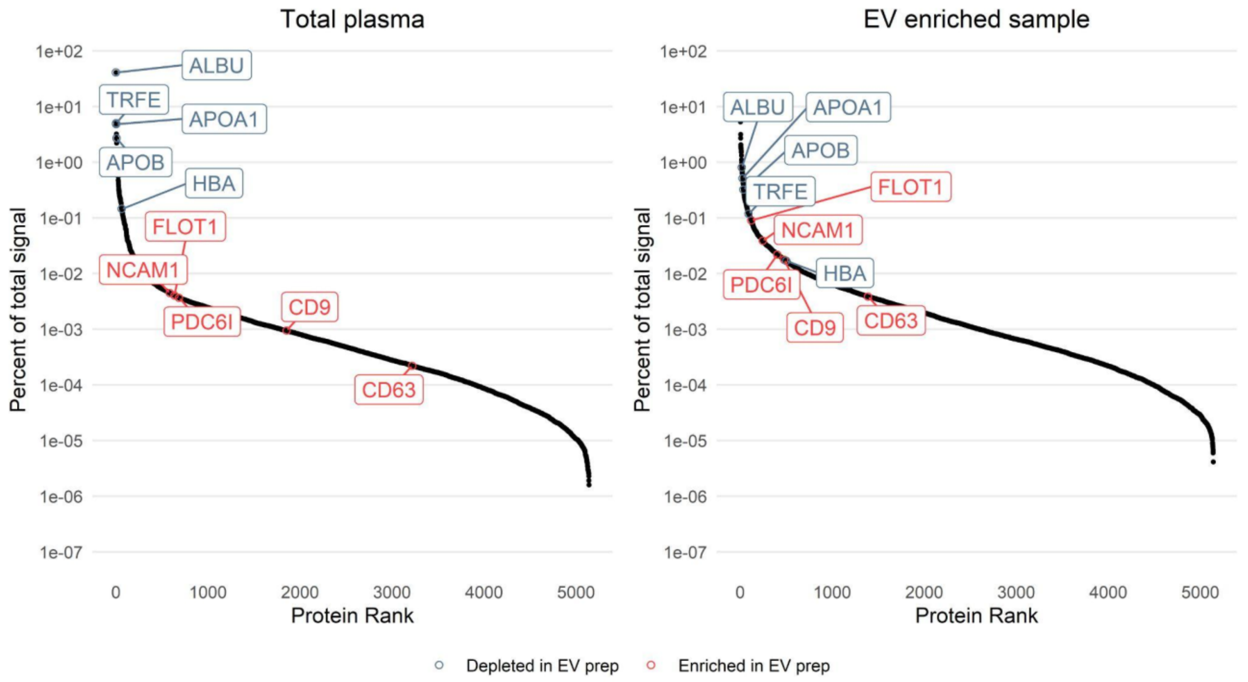


**Figure S16**. *Plasma dynamic range*. Abundance of plasma proteins detected in total plasma and EV enriched sample. Certain relevant marker proteins are annotated. Both plots show the same set of proteins, although many of them are essentially 0 in the total plasma sample as they are not detectable.

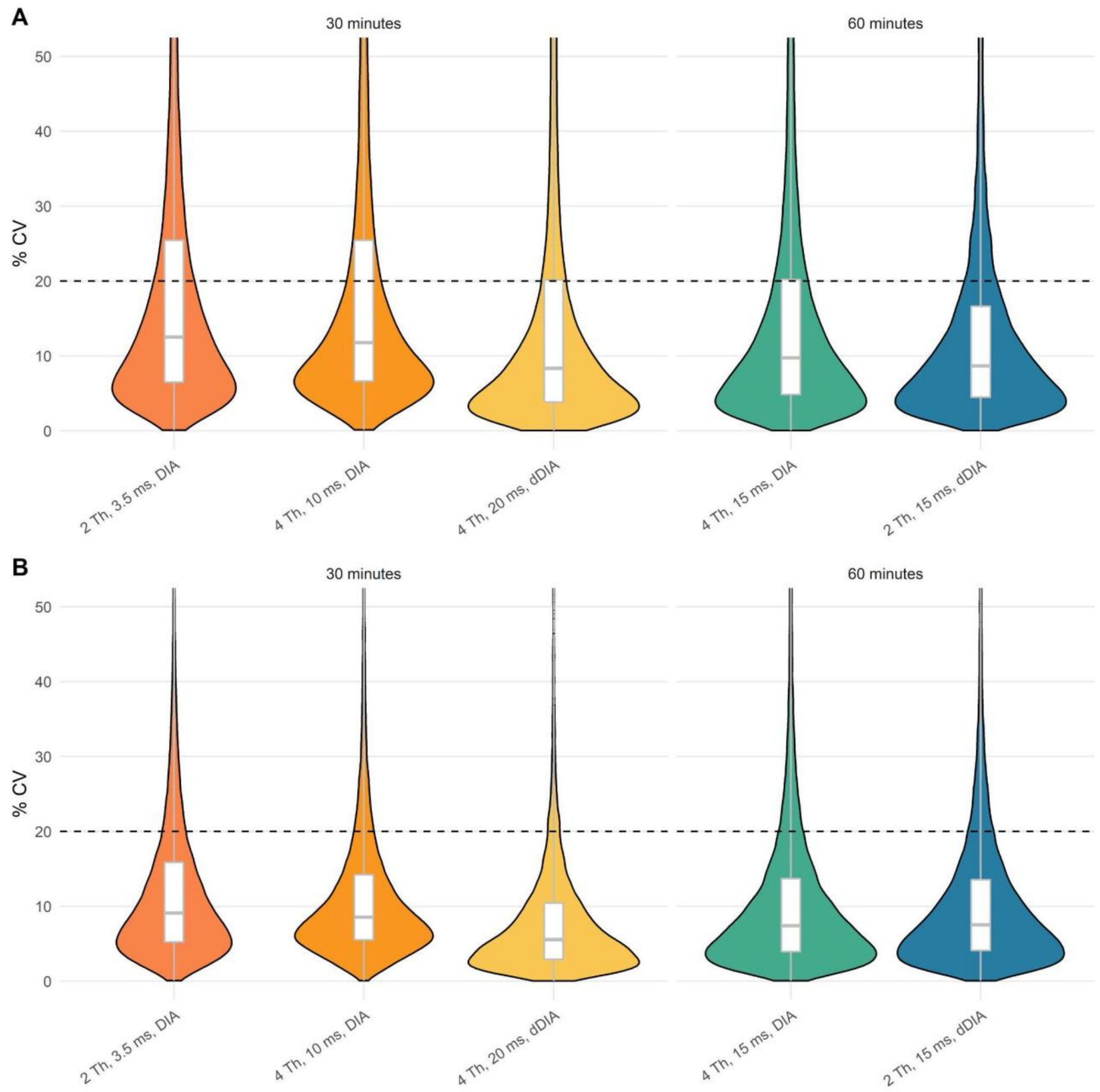


**Figure S17**. *Plasma CVs before and after refinement*. CVs before (A) and after (B) refinement for peptides with at least 3 co-eluting transitions. Note that the set of transitions was also filtered such that only the transitions that formed a co-eluting peak were integrated in the refined set. Black dashed line is 20% CV cutoff.
